# The mutability of demographic noise in microbial range expansions

**DOI:** 10.1101/2020.10.27.357483

**Authors:** QinQin Yu, Matti Gralka, Marie-Cécilia Duvernoy, Megan Sousa, Arbel Harpak, Oskar Hallatschek

## Abstract

Demographic noise, the change in the composition of a population due to random birth and death events, is an important driving force in evolution because it reduces the efficacy of natural selection. Demographic noise is typically thought to be set by the population size and the environment, but recent experiments with microbial range expansions have revealed substantial strain-level differences in demographic noise under the same growth conditions. Many genetic and phenotypic differences exist between strains; to what extent do single mutations change the strength of demographic noise? To investigate this question, we developed a high-throughput method for measuring demographic noise in colonies without the need for genetic manipulation. By applying this method to 191 randomly-selected single gene deletion strains from the *E. coli* Keio collection, we find that a typical single gene deletion mutation decreases demographic noise by 8% (maximal decrease: 81%). We find that the strength of demographic noise is an emergent trait at the population level that can be predicted by colony-level traits but not cell-level traits. The observed differences in demographic noise from single gene deletions can increase the establishment probability of beneficial mutations by almost an order of magnitude higher than the wild type. Our results show that single mutations can substantially alter adaptation through their effects on demographic noise and suggest that demographic noise can be an evolvable phenotype of a population.

## Introduction

Demographic noise, also referred to as “genetic drift”, “neutral drift”, or “drift”, is the change in the composition of a population due to random births and deaths. Theoretical population genetics predicts that demographic noise competes with natural selection by lowering the establishment probability of beneficial mutations (1) and causing the accumulation of deleterious mutations (2,3), leading to consequences such as the existence of a drift barrier (4) (a minimum absolute value fitness above which selection can act) and Mueller’s ratchet (5). Additionally, demographic noise reduces neutral genetic diversity (6), can limit mutation rates (7), and can also promote cooperation in spatially-structured environments (8). Experimental studies have validated many of these predictions (9–11), and demographic noise has been shown to play an important role in the evolutionary dynamics of a variety of systems including organelles (12), intestinal crypt stem cells (13), biofilms (14), the transmission of viruses (15–17) and human mitochondrial DNA (18), well-mixed culture (19), and potentially some types of cancer tumors (20).

Intuitively, the randomness of individual birth and death events should matter only relative to the population’s size (which be influenced by the environment), which is conventionally thought to set the strength of demographic noise (21–30). However, recent work in microbial colonies has shown that different strains from the same species can exhibit different strengths of demographic noise under the same growth conditions (26,31), and that the observed differences in demographic noise can have a substantial impact on the establishment probability of beneficial mutations (26,32,33). However, it is unknown how much single mutations can affect the strength of demographic noise and whether those changes would be sufficient to alter the efficacy of natural selection. In this work, we focus on loss of function mutations using single gene deletion mutant strains, as loss of function mutations are a common type of single step mutation in microbes.

Measuring the strength of demographic noise for a large number of strains requires a method for high-throughput tracking of cellular lineages in growing colonies. Previous methods for measuring demographic noise in microbial colonies required genetic transformations (31) or time-intensive microscopy and image analysis (26), which are impractical for testing a large number of strains. Here, we develop a label-free method to sparsely track cell lineages in growing colonies and use it to measure the distribution of demographic noise effects in *E. coli* single gene deletion strains. We show that most gene deletions decrease the strength of demographic noise, which in turn can dramatically increase the establishment probability of beneficial mutations. Our high-throughput approach also allows us to show that population-level emergent properties such as colony shape and size, but not single-cell properties such as cell shape, can predict the strength of demographic noise.

## Materials and Methods

### Strains and growth conditions

Single gene deletion strains were taken from the Keio collection (34), which consists of all non-essential single gene deletions in *E. coli* K-12 strain BW25113. Plasmids pQY10 and pQY11 were created by Gibson assembly of Venus YFP A206K (for pQY10) or Venus CFP A206K (for pQY11) (31), and Spec^R^ from pKDsgRNA-ack (gift from Kristala Prather, Addgene plasmid # 62654, http://n2t.net/addgene:62654; RRID:Addgene_62654) (35). Plasmids pQY12 and pQY13 were created similarly but additionally with and Cm^R^ from pACYC184.

All *E. coli* experiments were performed in LB (Merck 110285, Kenilworth, New Jersey) with the appropriate antibiotics and experiments with *S. cerevisae* were performed in YPD (36). All agar plates were prepared in OmniTrays (Nunc 242811, Roskilde, Denmark, 12.8cm×8.6cm) or 12cm×12cm square petri dishes (Greiner 688102, Kremsmuenster, Austria) filled with 70 mL media solidified with 2% Bacto Agar (BD 214010, Franklin Lakes, New Jersey). After solidifying, the plates were dried upside-down in the dark for 2 days and stored wrapped at 4C in the dark for 7-20 days before using.

### Fluorescent tracer beads

For experiments with *E. coli*, 1 μm red fluorescent polystyrene beads from Magsphere (PSF-001UM, Pasadena, CA, USA) were diluted to 3 μg/mL in molecular grade water and 920 μL was spread on the surface of the prepared OmniTray agar plates with sterile glass beads. Excess bead solution was poured out, and the plates were dried under the flow of a class II biosafety cabinet (Nuaire, NU-425-300ES, Plymouth, MN, USA) for 45 minutes. The bead density was chosen to achieve ~250 beads in a 56x field of view. For experiments with *S. cerervisiae,* 2μm dragon green fluorescent polystyrene beads from Bang’s labs (FSDG005, Fishers, IN, USA) were used at a similar surface density.

### Measurement of the distribution of demographic noise

We randomly selected 352 single gene deletion strains from the Keio collection. For each experiment, cells were thawed from glycerol stock (see **Supplementary methods**), mixed, and 5 μL was transferred into a 96-well flat bottom plate with 100μL LB and the appropriate antibiotics. Plates were covered with Breathe-Easy sealing membrane (Diversified Biotech BEM-1, Doylestown, PA, USA) and grown for 12 hours at 37C without shaking. A floating pin replicator (V&P Scientific, FP12, 2.36 mm pin diameter, San Diego, CA, USA) was used to inoculate a 2-3mm droplet from each well of the liquid culture onto a prepared OmniTray covered with fluorescent tracer beads. Droplets were dried and the plates were incubated upside down at 37C for 12 hours before timelapse imaging.

To account for systematic differences between plates, we also put 8 wild type BW25113 wells in each 96-well plate in different positions on each plate. The mean squared displacement (MSD, see below) of each gene deletion colony was normalized to the weighted average MSD of the wild type BW25113 colonies on that plate, ⟨MSD⟩_WT,_ and this “relative MSD” is reported. We performed three biological replicates for each strain (grown from the same glycerol stock), and their measurements were averaged together weighted by the inverse of the square of their individual error in relative MSD. The reported error for the strain is the standard error of the mean. During the experiment, several experimental challenges impede our ability to measure demographic noise, including the appearance of beneficial sectors (identified as diverging bead trajectories that correspond to bulges at the colony front) either due to de novo beneficial mutations or standing variation from glycerol stock (see **Supplementary section 2.4**), slow growth rate leading to bead tracks that were too short for analysis, no cells transferred during inoculation with our pinning tool, inaccurate particle tracking due to beads being too close together, or out of focus images. In order to keep only the highest quality data points, we focused on the 191 strains that had at least 2 replicates free of such issues.

### Timelapse imaging of fluorescent beads

Plates were transferred to an ibidi stagetop incubator (Catalog number 10918, Gräfelfing, Germany) set to 37C for imaging. Evaporation was minimized by putting wet Kim wipes in the chamber and sealing the chamber with tape. The fluorescent tracer beads at the front of the colony were imaged with a Zeiss Axio Zoom.V16 (Oberkochen, Germany) at 56x magnification. A custom macro program written using the Open Application Development for Zen software was used to find the initial focal position for each colony and adjust for deterministic focus drift over time due to slight evaporation. Timelapse imaging was performed at an interval of 10 minutes for 12 hours, during which time the colony grew about halfway across the field of view. Two z slices were taken for each colony and postprocessed to find the most in-focus image to adjust for additional focus drift. Subpixel-resolution particle tracking of the bead trajectory was achieved using a combination of particle image velocimetry and single particle tracking (37) and is described in detail in the **Supplementary methods**.

### Measurement of bead trajectory mean squared displacement

The measurement of mean squared displacement (MSD) is adapted from (31) and is illustrated in **Figure 1a** and **S1a**. Points in a trajectory that fall within a window of length *L* are fit to a line of best fit. The MSD is given by

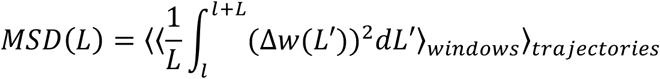

where *Δw(L’)* is the displacement of the bead trajectory from the line of best fit at each point, ⟨⟩_windows_ is an average over all possible definitions of a window with length *L* along the trajectory (window definitions are overlapping), and ⟨⟩_trajectories_ is a weighted average over all trajectories in a field of view, where the weight is the inverse squared standard error of the mean for each trajectory’s MSD(L) (**Figure S1a)**. We use 200 linearly spaced window sizes from *L* = 6 to 1152 μm. Window sizes that fit in fewer than 5 trajectories are dropped due to the noisiness in calculating the averaged MSD(L). Because we expect the trajectories to follow an anomalous random walk (31), the combined MSD(L) for all trajectories across the field of view is fit using weighted least squares to a power law, where the weight is the inverse square of the propagated standard error of the mean. Colonies with data in fewer than 5 window sizes are dropped due to the noisiness in fitting to a power law. The fit is extrapolated to L = 50 μm to give a single summary statistic for each colony, and this quantity is reported as MSD(L = 50 μm) (see **Supplementary section 2.2**), and the error is calculated as half the difference in MSD(L = 50μm) from using the upper and lower bounded coefficients to the fit.

**Figure 1:**
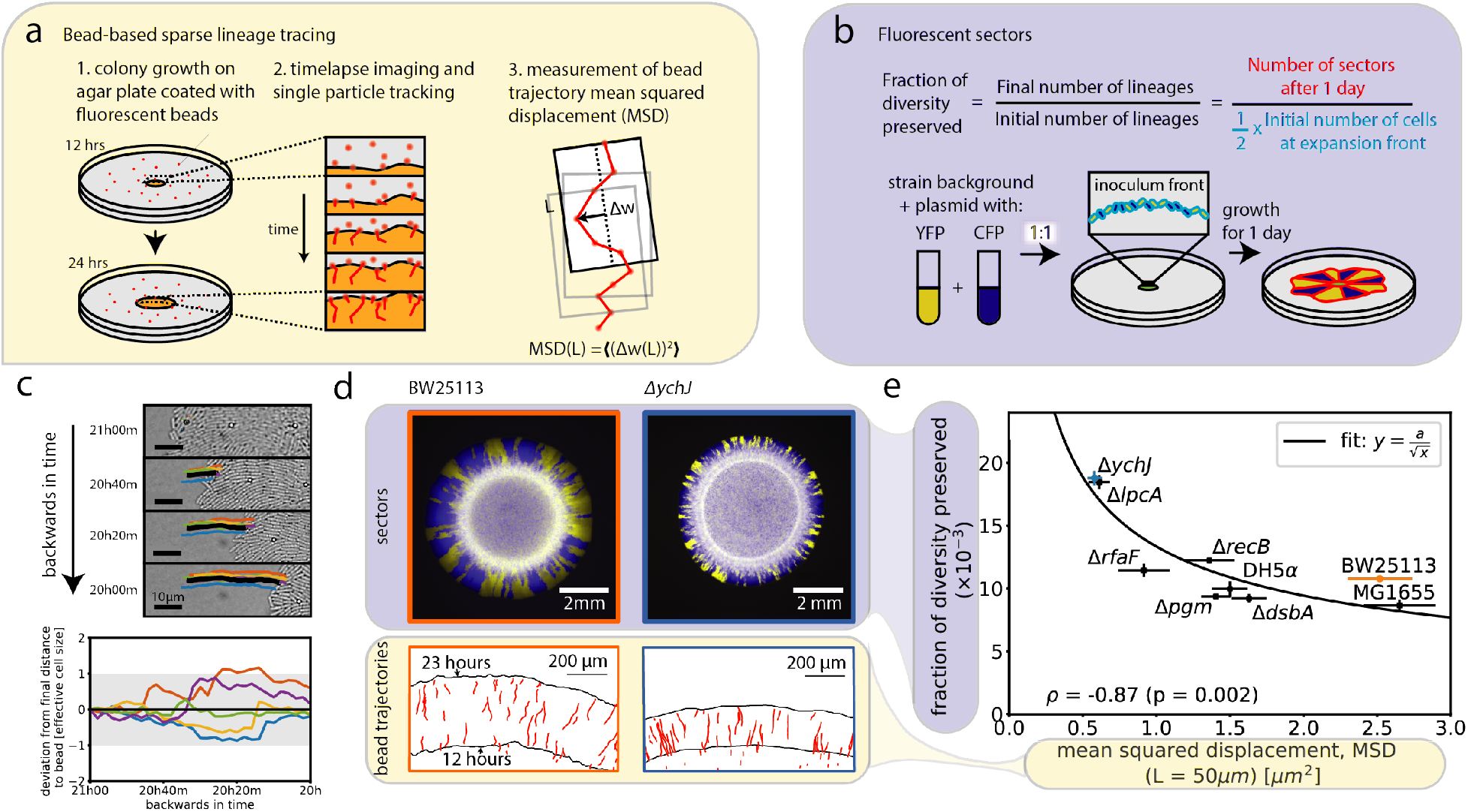
Label-free method of measuring demographic noise in microbial colonies. (a) Schematic of bead-based sparse lineage tracing method for measuring demographic noise. (b) Schematic of existing method for measuring fraction of diversity preserved (26). (c) (Top) The trajectory of a single bead (black) and the lineages of the cells neighboring it in the final-timepoint (colors) traced backwards in time in the Keio collection wild type strain. (Bottom) The deviation of the distance between the cell lineages and the bead from the final distance, backwards in time. Colors are the same as in the time series images. The gray shaded region shows a single cell width away or towards the bead. All cells that neighbor the bead in the final timepoint, except for one (orange), are neighbors of the bead in the first timepoint and stay within a single cell width of the final distance to the bead. (d) Example neutral mixtures of YFP and CFP tagged strains grown for 1 day and bead trajectories for strains highlighted in (e). (e) Comparison of MSD at window size *L* = 50 μm to the fraction of diversity preserved for 3 *E. coli* strain backgrounds and 6 single gene deletions on the Keio collection wild type background (BW25113). Error bars in MSD represent the standard error of the weighted mean (N = 7-8, see **Methods**) and error bars in the fraction of diversity preserved represent the standard error of the weighted mean (N = 8) where weights come from uncertainties in counting the number of sectors.

### Measurement of phenotypic traits

For the phenotypic trait measurements, in addition to the 191 single gene deletions, we also measured 41 additional strains of *E. coli* which included 4 strain backgrounds, 1 *mreB* knockout in the MC1000 background, 2 adhesin mutants, and 34 single gene knockouts from the Keio collection that we predicted may have large changes to demographic noise because of an altered biofilm forming ability in liquid culture (38) or altered cell shape from the wild type (using the classification on the Keio website, https://shigen.nig.ac.jp/ecoli/strain/resource/keioCollection/list). We normalized all phenotypic trait values to the average value measured from the wild type colonies on the same plate. The reported values for each strain are averages across 2-3 replicate colonies on different plates and the errors are the standard error of the mean. See the **Supplementary methods** for more details of the specific phenotypic trait measurements.

### Measurement of neutral fraction of diversity preserved

Neutral fluorescent pairs were created by transforming background strains with plasmids pQY10 (YFP, Spec^R^) or pQY11 (CFP, Spec^R^). Cells were streaked from glycerol stock and a single colony of each strain was inoculated into a 96 well plate with 600 μL LB and 120 μg/mL spectinomycin for plasmid retention. Plates were covered with Breathe-Easy sealing membrane and grown for 12 hours at 37C without shaking. 50 μL of culture from each strain in a neutral pair were mixed and a floating pin replicator was used to inoculate a 2-3mm droplet from the liquid culture onto a prepared OmniTray covered with fluorescent tracer beads. Droplets were dried and the plates were incubated at 37C.

Colonies were imaged after 24 hours with fluorescence microscopy using a Zeiss Axio Zoom.V16 and the number of sectors of each color was manually counted. The fraction of diversity preserved was calculated as in Ref (26) by dividing the number of neutral sectors by one-half times the estimated initial number of cells at the inoculum front (see **Figure 1b**). The factor of one-half accounts for the probability that two neighboring cells at the inoculum front share the same color label. The initial number of cells is estimated by measuring the inoculum size of each colony (manually measured by fitting a circle to a brightfield backlight image at the time of inoculation) divided by the effective cell size for *E. coli* (taken to be 1.7 μm, Ref (26)).

### Colony fitness

The colony fitness coefficient between two strains was measured using a colony collision assay as described in Refs (26,39) by growing colonies next to one another and measuring the curvature of the intersecting arc upon collision. Cells were streaked from glycerol stock and a single colony for each strain was inoculated into LB with 120 μg/mL spectinomycin for plasmid retention and incubated at 37C for 15 hours. The culture was back diluted 1:500 in 1mL fresh LB with 120 μg/mL spectinomycin and grown at 37C for 4 hours. 1 μL of the culture was then inoculated onto the prepared 12cm×12cm square petri dishes containing LB with different concentrations of chloramphenicol (0μg/mL, 1μg/mL, 2μg/mL, 3 μg/mL) in pairs that were 5 mm apart, with 32 pairs per plate, then the colonies were incubated at 37C. After half of a day, bright field backlight images are taken and were used to fit circles to each colony to determine the distance between the two colonies. After 6 days, the colonies were imaged with fluorescence microscopy using a Zeiss Axio Zoom.V16. The radius of curvature of the intersecting arc between the two colonies was determined with image segmentation and was used to calculate the fitness coefficient between the two strains (**Figure S14a**).

### Measurement of non-neutral establishment probability

We transformed 9 gene deletion strains from the Keio collection (gpmI, recB, pgm, tolQ, ychJ, lpcA, dsbA, rfaF, tatB) and 3 strain backgrounds (BW25113, MG1655, DH5⍺) with pQY11 (CFP, Spec^R^) or pQY12 (YFP, Spec^R^, Cm^R^). Cells were streaked from glycerol stock and a single colony of each strain was inoculated into media with 120 μg/mL spectinomycin for plasmid retention, then incubated at 37C for 16 hours. The culture was back-diluted 1:1000 in 1mL fresh media with 120 μg/mL spectinomycin and grown at 37C for 4 hours. YFP chloramphenicol-resistant and CFP chloramphenicol-sensitive cells from the same strain background were mixed respectively at approximately 1:500, 1:200, and 1:50 and distributed in a 96-well plate (**Figure 4a**). A floating pin replicator was used to inoculate a 2-3mm droplet from the liquid culture onto prepared OmniTrays with varying concentrations of chloramphenicol (0μg/mL, 1μg/mL, 2μg/mL, 3 μg/mL). Droplets were dried and the plates were incubated at 37C for 3 days, then imaged by fluorescence microscopy using a Zeiss Axio Zoom.V16.

The establishment probability of the beneficial strain is measured as described in (26). Briefly,

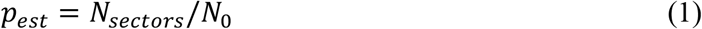

where *N_sectors_* is the number of resistant sectors after 3 days (counted by eye) and *N_0_* is the estimated initial number of cells of the resistant type at the inoculum front. Because the establishment probability can only be accurately measured when the initial number of resistant cells is low enough that the resistant sectors do not interact with one another, we only keep colonies where neighboring resistant sectors are distinguishable at the colony front. The initial number *N_0_* of cells of the resistant type is estimated by multiplying the initial number of cells at the inoculum front (see measurement of neutral fraction of diversity preserved) by the fraction of resistant cells in the inoculum (measured by plating and counting CFUs).

## Results

### Label-free method for measuring demographic noise in microbial colonies

To measure demographic noise in an expanding microbial colony without genetic labels, we developed a method that consists of two steps (**Figure 1a**, **Methods,** and **Supplementary section 2.1**). First, we record the trajectories of cell-sized fluorescent beads embedded in the colony (**Supplementary movie**), which track lineages of their neighboring cells (**Figure 1c**, **Figure S1c-d,** and **Supplementary methods**). Second, we analyze the fluctuations of the measured lineages (via the bead trajectories) using their length-dependent mean squared displacement (MSD), which serves as an established statistic to quantify the strength of demographic noise (**Methods** and Ref (31)).

To determine the ability of our method to measure differences in demographic noise, we compared it to an existing method which uses neutral fluorescent labels to measure the fraction of diversity preserved (the fraction of surviving fluorescent sectors) after a range expansion (31,40) (**Figure 1b** and **Methods**). **Figure 1d** shows that the fraction of diversity preserved is negatively correlated with MSD for a subset of 9 strains (*ρ* = −0.87, p = 0.002), and can be well fit to an inverse square root relationship 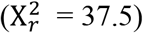. This inverse square root relationship is consistent with the theory expectation for the fluorescent sector boundary MSD (40), suggesting that the bead MSD captures the fluorescent sector boundary MSD, and is thus a convenient and reliable measure of demographic noise. We chose to report the MSD at the window length *L* = 50μm because the inverse square root fit to the fraction of diversity preserved had the lowest chi-squared at this length scale (**Figure S3**). To control for growth rate differences between the strains, we also masked the colonies with the smallest colony’s outline and remeasured the fraction of diversity of preserved; this did not significantly change the ordering of the genotypes (**Figure S1b**).

### The distribution of demographic noise for single gene deletions

We next wanted to use our bead-based sparse lineage tracing method to measure the distribution of demographic noise due to single gene deletion mutations. We randomly selected 191 single gene deletion strains from the Keio collection (34), a well-characterized library of *E. coli* strains that contains all non-lethal single gene deletions (see **Methods**). In order to test such a large number of colonies, we grew the colonies in 96-array format on multiple agar plates. We observed variation in bead MSD between plates (see **Supplementary section 2.3**), and therefore report the bead MSD of the gene deletion strains relative to that of the wild type, which is present in 8 replicate colonies per plate (see **Methods**).

**Figure 2** shows the distribution of relative MSD from the 191 randomly selected single gene deletion strains. The knockout (KO) distribution significantly differs from the wild type (WT) distribution (Kolmogorov-Smirnov p = 2.7×10^−4^) with a lower mean (KO: 0.904, WT: 1.011) and higher variance (KO: 0.044, WT: 0.004). 39% of knockout MSDs were lower than the lowest wild type MSD observed, with the maximal decrease in knockout MSD of 81% from the wild type median MSD. Interestingly, the typical knockout mutation decreases demographic noise from that of the wild type by 8% (95% CI = [1%, 17%], **Supplementary Methods**) (Median KO = 0.94, Median WT = 1.02).

**Figure 2:**
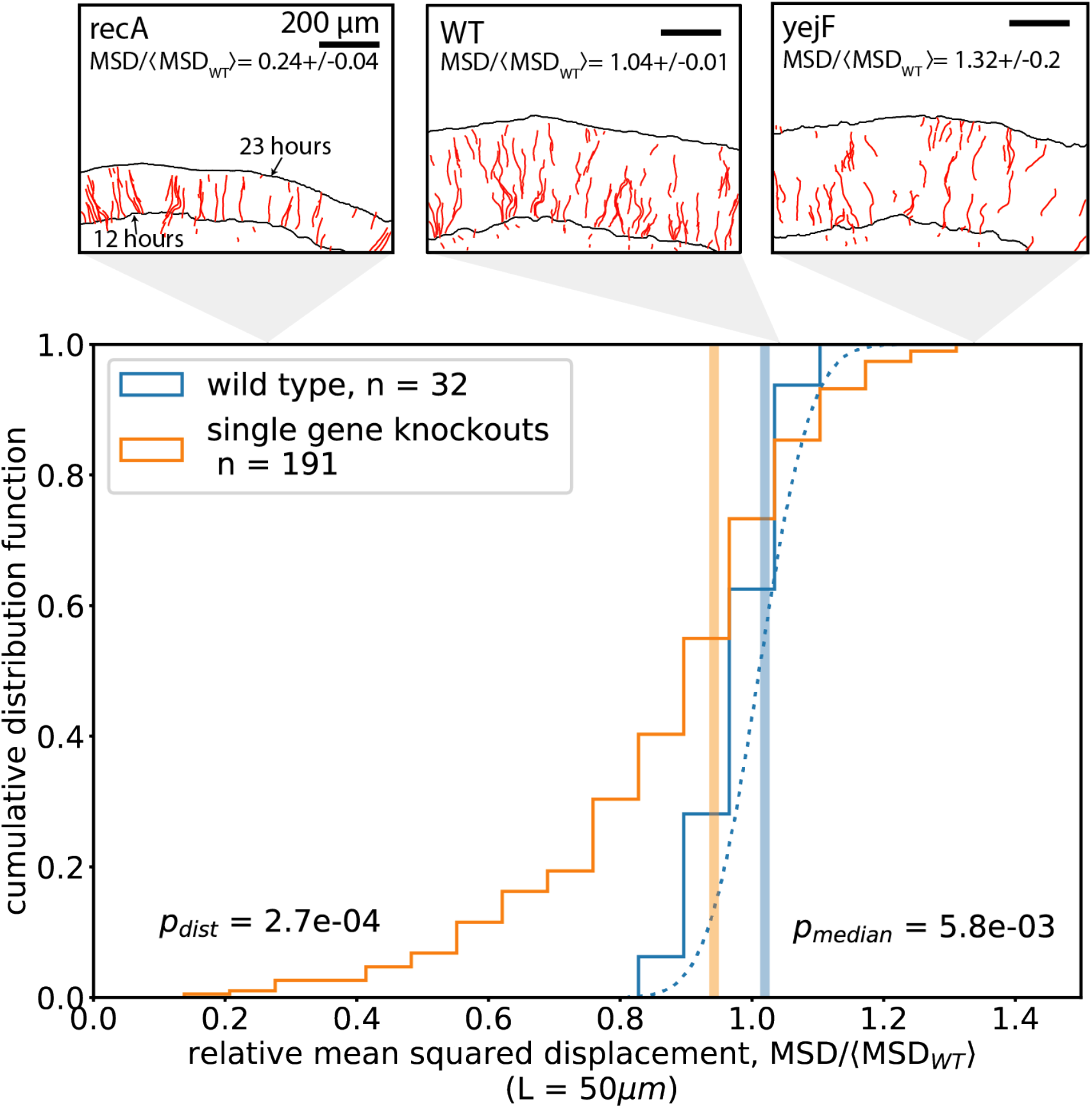
Distribution of MSD across randomly-sampled Keio collection single gene deletion strains. Each count in the distributions is an average of 2-3 replicate colonies grown on different agar plates, and the MSD is normalized to the average wild type MSD on each plate (**Methods**). The blue dotted line shows a Gaussian fit to the wild type distribution. Vertical lines show the median value of each distribution. Panels show examples of bead trajectories from wild type and single gene deletions strains from the Keio collection. Black lines in panels show the colony front at t = 12 hours and t = 23 hours.

### Phenotypic trait predictors of the strength of demographic noise

We noticed that some colonies with particularly low bead MSD also seemed to be small with smooth colony shapes. As a result, we systematically checked which phenotypic traits best correlate with the observed differences in strengths of demographic noise in the single gene deletions. Specifically, we measured a range of traits in the same colonies for which we measured bead MSD (see **Methods**), including the depth of the growing layer of cells at the front, the roughness of the colony front, the area of colony, and we also used existing datasets for single cell shape. While previous studies have studied the relationship of these traits with demographic noise experimentally by comparing species or strains in low-throughput (26,32,41), the same strain in different nutrient concentrations (27), or using simulations (8,32), our system allows us to experimentally test correlations of demographic noise with different traits in a large number of related strains, thus overcoming a major experimental limitation.

We found that (1) the roughness of the colony front is positively correlated with the bead MSD (**Figure 3a**, Pearson *r* = 0.66, p = 2×10^−23^), (2) the size of the growing layer of cells at the front of the colony is negatively correlated with the bead MSD (**Figure 3b**, Pearson *r* = −0.54, p = 4×10^−15^), and (3) the colony area after 1 day of growth, which we checked can be used as a proxy for biomass (see **Supplementary methods** and **Figure S9a**-**b**), is positively correlated with the MSD (**Figure 3c**, Pearson’s ⍴ = 0.63, p = 5×10^−21^). These colony-level results agree with theoretical predictions (8,32) and previous experimental results (27,32). Using datasets of single cell shapes from the Keio collection from Refs (42,43), we did not find a significant correlation of demographic noise with cell shape (**Figure S8a**-**b**), in contrast to the colony-level traits.

**Figure 3:**
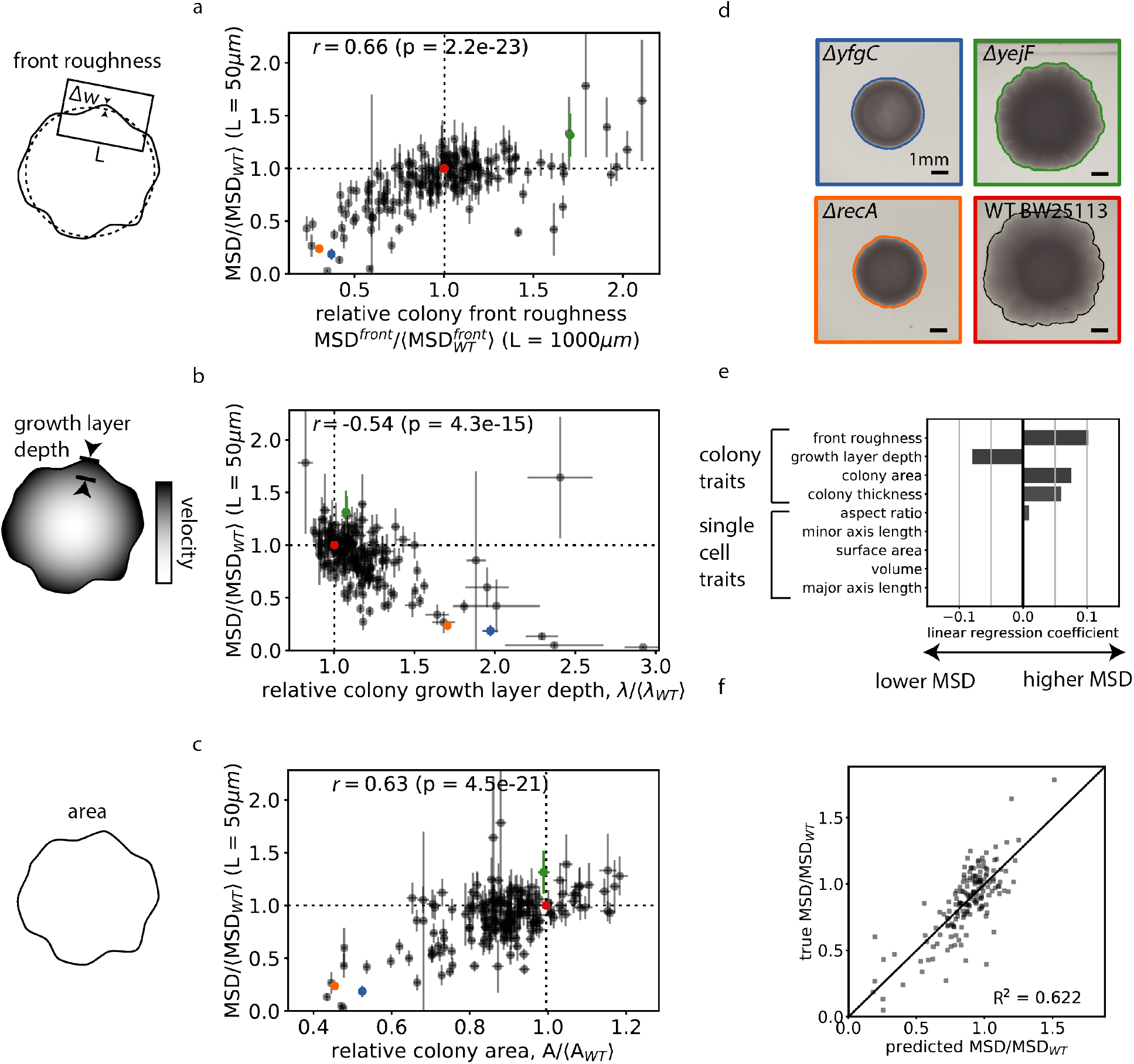
Phenotypic predictors of the strength of demographic noise. Correlation of the bead trajectory MSD for 191 single gene deletions and 41 selected strains with (a) front roughness (defined in **Supplementary methods**), (b) colony growth layer depth (defined in **Supplementary methods** and **Figure S9c-d**), and (c) colony area. Error bars represent the standard error of the mean across 2-3 replicate colonies. (d) Example colonies for colored points in (a)-(c) (e) Linear model coefficients for phenotypic traits that best predict MSD, estimated through Lasso regression. (f) Predicted MSD using the linear model with the coefficients shown in (e).

We estimated the joint relationship of the measured traits with bead MSD using Lasso regression (44), which finds the minimal set of traits that predict the MSD and the coefficients associated with those traits in a linear model. The traits that were included in the Lasso regression were the 4 colony traits (front roughness, growth layer depth, colony area, and colony thickness) that we measured and the 5 single-cell shape traits (aspect ratio, minor axis length, surface area, volume, and major axis length) from the dataset in Ref (43). We find that all 4 colony level traits and single cell aspect ratio are the only 5 traits included in the best fit model to the MSD (**Figure 3e**), with the coefficient for the single cell aspect ratio being almost an order of magnitude lower than that of the lowest colony trait coefficient. Using the best fit model, we are able to explain the variance in the MSD with an R^2^ of 62% (**Figure 3f**).

These correlations of demographic noise with various population-level traits could be partly driven by correlations between the population-level traits themselves. Indeed, we find that colonies with larger areas tend to have smaller growth layer depths (**Figure S10b**) and higher front roughness (**Figures S10a**). Prior theoretical studies have suggested that colony traits are interdependent (8,32): faster growing strains have a sharper nutrient gradient at the front, leading to a smaller growth layer, which in turn creates more front roughness, which is consistent with our findings. Both the correlation of colony area with front roughness and that of colony area with bead MSD across strains could potentially be explained without demographic noise differences if the front roughness and bead MSD increased over time within a single colony as it grew larger. In order to exclude this possibility, we checked that the front roughness saturates over time by the time of measurement (**Figure S9f**) and that the bead MSD does not increase as the colony grows larger but rather slightly decreases (**Figure S9e)**. In order to test which traits can be a causal determinant of MSD, we corrected for linked correlations between traits using partial correlations (**Figures S11**), and find a slightly lower but significantly nonzero correlation of bead MSD with front roughness (r = 0.53, p = 2e-13) and a more substantial decrease in correlation of bead MSD with colony area (r = 0.29, p = 2e-4) and growth layer depth (p = −0.17, p = 2.5e-2). This supports the idea that front roughness is the main causal determinant of MSD, as was also shown in Ref (32).

Because previous work has found that colonies grown from cells with round shapes tend to have lower demographic noise (26,41), we were puzzled by our result showing lack of correlation between cell shape and bead MSD. Thus, we specifically tested a round cell shape mutant, MC1000 *ΔmreB*, which we indeed measured to have low MSD compared to the wild type MC1000 (**Figure 8e**). However, by using the best fit Lasso regression model (**Figure 3e**), which primarily includes colony-level traits, the low MSD could be predicted (**Figure S8e**), suggesting that colony-level traits are sufficient to explain the difference in MSD. Because it is possible that differences in colony-level traits mask the effect of single-cell traits on bead MSD, we also corrected for variation in all other traits using partial correlations; however, the corrected bead MSD still shows little correlation between cell shape and bead MSD (**Figure S8f**). We note that we cannot rule out the possibility that the lack of correlation between bead MSD and cell shape for the Keio mutants described above using the datasets from Refs (43,45) is influenced by differences in cell shape exhibited by cells of the same genotype in different growth conditions (see **Figure S8c-d** and **Supplementary section 2.7**). We also measured a library of *mreB* and *mrdA* point mutants that were enriched for cell shape differences (45). In this enriched library, there is a slightly higher correlation between cell shape and the strength of demographic noise (Pearson *r* =0.32 and *r* = 0.46 for *mreB* and *mrdA* point mutants respectively, see **Figures S8g-h**), possibly because the cell shapes span a larger range, or because our growth condition was more similar to that of the single cell measurements in this dataset.

In summary, Lasso regression suggests that a combination of colony-level traits best predicts bead MSD, which we have shown is anticorrelated with demographic noise. However, after correcting for correlations between phenotypic traits, we found evidence supporting that the main causal determinant of MSD is the colony front roughness. Additionally, the agreement of the colony-level phenotypic trait relationships with those found in previous work (8,26,27,32) suggests that the same mechanisms for how phenotypes affect demographic noise seem to hold in range-expanding populations regardless of whether looking across single mutations, different strains, or different species.

### Single gene deletions can substantially alter adaptation through changes to demographic noise

Finally, we sought to determine whether the variation in demographic noise induced by single deletions also induces a substantial corresponding change in evolutionary outcomes, such as the establishment probability of beneficial mutations, as predicted by population genetics theory. We constructed fluorescently-labeled chloramphenicol resistant and sensitive strains on selected strain backgrounds and measured the establishment probability of the resistant type when competed with the sensitive type on the same strain background (**Figure 4a** and **Methods**). The fitness coefficient between the resistant and sensitive types was tuned by the chloramphenicol concentration and measured for each strain background at each chloramphenicol concentration using a colony collision assay (**Methods** and **Figure S14a**).

**Figure 4:**
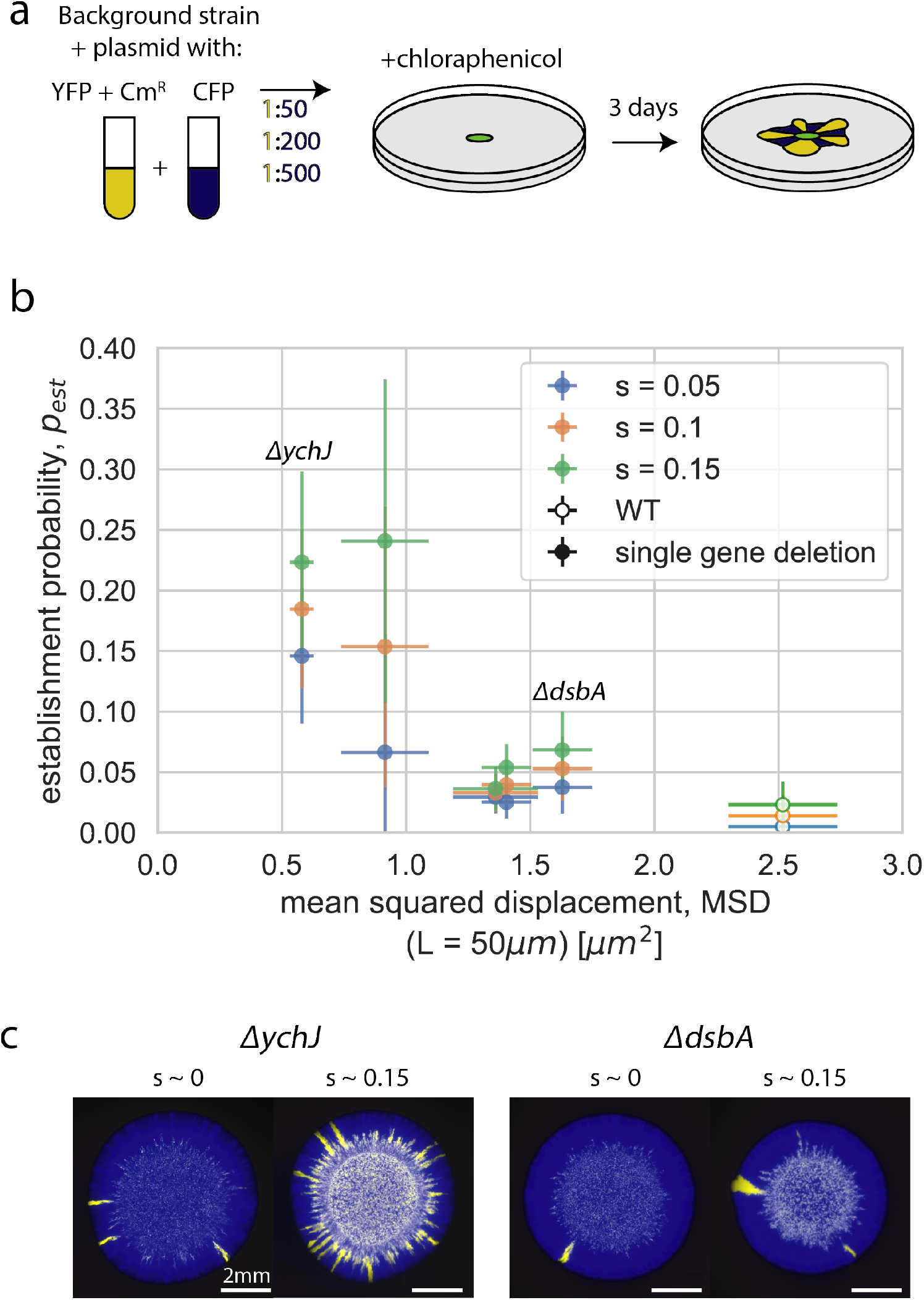
Single gene deletions can substantially alter adaptation through changes to demographic noise. (a) Schematic for measurement of establishment probability. After 3 days, the number of beneficial sectors is counted and the establishment probability is calculated by dividing by the initial number of chloramphenicol resistant cells at the expansion front (26). Cm^R^: chloramphenicol resistance gene. (b) Interpolated establishment probability at three different fitness coefficients as a function of bead trajectory MSD. Error bars in establishment probability represent linear fitting error (see **Supplementary methods**) and error bars in bead MSD represent the standard error of the weighted mean (N = 7-8, see **Methods**) (c) Example colonies for two different strain backgrounds each at two fitness coefficients between the resistant and sensitive types.

We found that the establishment probability of the resistant type is negatively correlated with the bead MSD of the strain background across beneficial fitness coefficients from s = 0.05 to s = 0.15 (**Figure 4b**). The Keio collection WT had the largest MSD and lowest establishment probability of the Keio collection strains that were tested. The maximal increase in the establishment probability of a beneficial mutant on a gene deletion background was about 6-fold over the WT, which corresponded to about a 4-fold decrease in the background strain MSD compared to the WT. Interestingly, we observed that that the establishment probability was decreased when the initial fraction of the resistant type was increased (**Figure S13**), possibly due to interactions between beneficial sectors; we controlled for differences in initial fraction by separating the data by initial fraction, and found that the effect of the initial fraction of the resistant type on the establishment probability does not explain the observed negative correlation between demographic noise and establishment probability (see **Figure S15b**). In sum, we find that the range of strengths of demographic noise accessible by single gene deletion strains substantially affects the establishment probability of a beneficial mutant on that background.

## Discussion

We have shown that single gene deletions can substantially alter the strength of demographic noise in microbial colonies (**Figure 2**) and that these differences can have an impact on adaptation (**Figure 4**). We accomplished this by developing a bead-based sparse lineage tracing method for measuring demographic noise in colonies (**Figure 1**). While the beads could potentially perturb the cell lineages, possibly impacting the correlation of bead MSD with the fraction of diversity preserved, both quantities allowed us to observe sufficient differences between strains. We checked whether there were particular types of genes that altered demographic noise but found no clear functional signatures that were enriched for (see **Supplementary section 2.5**). We additionally used this method to measure a non-random set of strains from the Keio collection as well as mreB and mrdA point mutants and found an even larger range of demographic noise effects (**Supplementary section 2.6** and **Figure S7**).

Our results suggest that demographic noise itself may be an evolvable trait of a population. We hypothesize that strain backgrounds with different strengths of demographic noise may also exhibit different rates of adaptation when accumulating multiple mutations. It would be interesting to test this hypothesis in future work empirically through experimental evolution of colonies (46,47) and theoretically through simulations with joint distributions of demographic noise effects and fitness effects. Quantitatively, the evolution of demographic noise may be similar to the evolution of mutation rate, because both mutations and demographic noise primarily influence the establishment rate of new mutations; however, this should be examined more carefully in future work in different regimes such as successive mutations and clonal interference (48). Additionally, interesting dynamics could arise in spatially-structured communities with cooperation, such as those that share a common good (8). Increasing demographic noise in these systems may make cheating less likely by leading to more spatial segregation of cheater and producer types. Another interesting corollary to our results is that a decrease in the strength of demographic noise enables more efficient transfer of genetic material through conjugation in bacterial colonies (49) and exchange of metabolites between co-expanding strains (50,51).

Our results show that demographic noise is correlated with colony-level traits (**Figure 3**), suggesting that the strength of demographic noise in these colonies is set by collective behavior. As a result, we hypothesize that the plasticity of demographic noise holds more generally in self-organized systems (52), including colonies, biofilms, spatially-structured microbiomes, and solid cancer tumors, which would be interesting avenues for future study. Additionally, other phenotypic traits have been predicted to influence colony patterning and demographic noise and it would be interesting to test their influence on demographic noise in future work, including that of cell-cell and cell-substrate adhesion (53–55), cell orientations (32), cell elasticity (32), and variation in single cell growth rates (56,57) and lag times (58,59).

The positive correlation between colony area and demographic noise (**Figure 3c**, r = 0.63) suggests a tradeoff between demographic noise and fitness (**Figure S12**): a beneficial mutation may increase demographic noise and actually impair its own establishment and once established, also the establishment of future beneficial mutations. However, when a demographic-noise-modifying mutant first arises at a low frequency in a colony, the colony-level traits will be set by both that of the mutant and the background strain, so the strength of demographic noise that governs its trajectory will likely be a complex time-dependent combination of traits from the two genotypes in monoculture. Thus, while this work generates interesting hypotheses as to the tradeoffs between demographic noise and fitness, future work is needed to more closely examine the consequences of demographic noise and fitness correlations in different environments.

The bead-based sparse lineage tracing method in colonies can be extended to study demographic noise in other genotypes in high-throughput, such as double-mutants and potentially other species. In the supplementary text, we use this method to measure demographic noise in *S. cerevisiae* colonies (**Figure S1e**), and we find a lower bead MSD in *S. cerevisiae* compared with that of *E. coli*, in agreement with results from previous work (26,31). Measurements of demographic noise in additional genotypes can be used to understand the dependence of the distribution of strength of demographic noise on the genetic background across large mutational differences (different species) or small mutational differences (double-mutants).

Like selection, mutation, migration, and recombination, demographic noise has been shown to be an important evolutionary force in many systems. Understanding the environmental and genetic influences on demographic noise will allow us to better identify and model the relevant forces that drive evolution in different systems. Whereas demographic noise is typically thought of as being static or dependent on the environment, we have shown that like for other evolutionary forces, demographic noise can be considered an evolvable trait of a population. Future work exploring the evolvability of demographic noise will help us better understand its consequences on evolutionary outcomes in different systems.

## Acknowledgements

We thank Jona Kayser, Joao Ascensao, Benjamin Good, and the Hallatschek lab for valuable discussions and feedback. We thank KC Huang and Handuo Shi for sharing the mreB and mrdA point mutation libraries, and Christophe Beloin for sharing the quandruple adhesin and polysaccharide mutant strains. We thank the Laura Waller lab for lending the objective used for cell shape validation. Research reported in this publication was supported by the National Institute of General Medical Sciences of the National Institutes of Health under award R01GM115851 to OH, a National Science Foundation CAREER Award (#1555330) to OH and a Simons Investigator award from the Simons Foundation (#327934) to OH, a National Science Foundation Graduate Research Fellowship under Grant No. (DGE 1106400) to QY, and a fellowship from the Simons Foundation’s Society of Fellows (#633313) to AH.

