## Supplementary text for "The mutability of demographic noise in microbial range expansions"

#### 1 Supplementary methods

##### 1.1 Strains

| Strain | Description | Genotype | Derived from | Antibiotics | Ref |
|---|---|---|---|---|---|
| BW25113 | Keio collection background | | *E. coli* K-12 | | [2] |
| BW25113 single gene deletions | Keio collection | see Supplementary Table 1 | BW25113 | | [2] |
| MG1655 | | | *E. coli* K-12 | | |
| DH5$\alpha$ | | | | | |
| MC1000 | | | | | [11] |
| MC1000 $\Delta mreB$ | | | MC1000 | | [11] |
| $\Delta_4$pol | Deletion mutant of polysaccharides Yjb, cellulose, PGA and colonic acid | MG1655 $\Delta yjbEH :: cm$ $\Delta bcsA :: Km$FRT $pgaA :: uidA - zeo$ $cps5$ ::Tn10 | | | [4] |

| | | | | | |
|---|---|---|---|---|---|
| $\Delta_4$adh | Deletion mutant of flagella, AG43, type 1 fimbriae, and curli | MG1655 $gfp$ $\Delta fliER$ :: $cm$ $\Delta fimAH$ :: $zeo$ $\Delta flu$ ::FRT $\Delta csgA$ :: $spec$ | | | [5] |
| RDM 893 | *mreB* point mutant background | MG1655 $\Delta mreB$ pRMmreBCD | | | [15] |
| TKL 117 | *mrdA* point mutant background | MG1655 $\Delta mrdA$ pRMind-pbp2 | | | [15] |
| *mreB* point mutants | | see Supplementary Table 2 | RDM 893 | 15 $\mu$g/mL chloramphenicol | [15] |
| *mrdA* point mutants | | see Supplementary Table 2 | TKL 117 | 35 $\mu$g/mL kanamycin and 50 $\mu$M IPTG | [15] |

#### 1.2 Plasmids

| Plasmid | Description | Genotype | Derived from | Antibiotics | Ref |
|---|---|---|---|---|---|
| pQY10 | | Venus YFP (A 206K), Spec$^R$ | | 120 $\mu$g/mL spectinomycin | this study |
| pQY11 | | (e)CFP (A 206K) , Spec$^R$ | | 120 $\mu$g/mL spectinomycin | this study |
| pQY12 | | Venus YFP (A 206K), Spec$^R$, Cm$^R$ | | 120 $\mu$g/mL spectinomycin, variable chlorampenicol | this study |

| | | | | | |
|---|---|---|---|---|---|
| pQY13 | | (e)CFP (A 206K) , Spec$^R$, Cm$^R$ | | 120 $\mu$g/mL spectinomycin, variable chlorampenicol | this study |

#### 1.3 Creation of glycerol stocks for measurement of distribution of demographic noise

We created master stocks by rearraying strains from the original frozen Keio collection glycerol stocks into 96-well plates with LB, growing overnight, and freezing the rearrayed cultures in a 25% glycerol stock. Master stocks of the mreB and mrdA point mutant libraries [15] were acquired from the lab of KC Huang. We created separate glycerol stocks that could be defrosted for each experiment by scraping frozen glycerol stock from the master stocks into a 96 deep well plate filled with 1 mL LB with the appropriate antibiotics, growing overnight until saturation, aliquoting 100 $\mu$L into multiple 96-well plates, and freezing as separate glycerol stocks.

#### 1.4 Single particle tracking

Single particle tracking was performed using a custom-written code in MATLAB based on that used in Ref [4]. Briefly, particle image velocimetry (PIV) was first used to subtract large-scale movements between frames. Then, particles with a radius smaller than 3 pixels were detected and linked between consecutive frames with single particle tracking to sub-pixel accuracy. Displacements between consecutive timepoints were then joined to create trajectories spanning multiple timepoints. The trajectories of particles in regions without cells were used to calibrate xy fluctuations in stage position over time. After correcting for stage fluctuations, trajectories that were shorter than 30 $\mu$m in total length were rejected as stationary beads and consecutive timepoint steps that were shorter than 1 pixel were rejected as noise.

#### 1.5 Measurement of colony front roughness

The front roughness was measured using the method described in Ref [7]. A backlight image of the colony was taken at 24 hours, and a custom-written MATLAB code was used to extract the colony boundary using image segmentation. The boundary was fit with a circle and the mean squared displacement in the radial direction was calculated for windows of different arc lengths ($L = 200$ linearly spaced arc lengths from 6 to 1152 $\mu$m) along the best fit circle using a running average over overlapping definitions of the starting position of the window. The MSD was fit to a power law as a function of $L$ and the value of the fit at $L = 1000\mu$m was reported.

#### 1.6 Measurement of growth layer depth

The growth layer depth was measured using bead displacements between pairs of consecutive timepoints (Figure S9c). We assumed that the velocity $v$ along the direction of growth has the form

$$v_y(y) = v_{y,max}e^{-(y-y_0)/\lambda} \tag{1}$$

where $v_{y,max}$ is the maximum velocity, $y$ is the position along the direction of growth, $y_0$ is the front position, and $\lambda$ is the growth layer depth. The ratio $v$ at two positions within the colony then conveniently takes on the form

$$v = \frac{v_y(y_2)}{v_y(y_1)} = e^{-(y_2-y_1)/\lambda}. \tag{2}$$

The average direction of motion of all the trajectories in a field of view was determined, and all subsequent measurements were projected along this axis. For each pair of consecutive timepoints, and for each pair of beads, the ratio of their displacements, $\frac{v_y(y_2)}{v_y(y_1)}$, and the distance between them parallel to the average direction of motion, $\Delta y = y_2 - y_1$, were calculated. Because noise from inaccurate tracking can dominate the ratio of the velocity when one or both of the bead velocities is small, we restricted our analysis to $-50\mu m < \Delta y < 50\mu m$. To minimize the effect of changes in curvature of the front of the colony, we restricted pairs of beads to have $\Delta x < 50\mu m$. We binned the data (bin size $= 10\mu m$, or if range of $\Delta y < 100\mu m$ then we used a bin size of std$(\Delta y)/2$) and fit to an exponential decay function using weighted least squares to extract the growth layer depth $\lambda$ (Figure S9d). The error in $\lambda$ represents the error in the fitting parameter.

#### 1.7 Measurement of colony thickness

The thickness of the colony was determined using a backlight brightfield image of the whole colony at 23h, where the exposure time was fixed for all colonies. We calibrated the intensity of the transmitted light to the colony thickness using a colony that was grown with fluorescent tracer beads (Methods in main text). We noticed that beads primarily rise to the top of the colony, so we used the height of the fluorescent beads as the ground truth colony height. To measure the average thickness of the whole colony, we calculated the average brightfield intensity within each colony and converted it into a thickness using the calibration curve. We note that this method does not distinguish between changes to intensity due to changes in colony thickness and changes in biomass density, but we make the assumption that any changes to density are small in comparison to changes in thickness.

#### 1.8 Single cell shape comparison in liquid and colony conditions

Cell preparation for growth in liquid culture was closely matched to the protocols of the National BioResource Project described in Refs [16] and [6]. Cultures were grown overnight with rotation at 30C, back-diluted 1:100 in fresh LB medium the next day and then incubated at 37C for 2 hours. Before imaging, cells were diluted in PBS to achieve the desired imaging density and vortexed to break up clumps. For imaging, a droplet of the diluted culture was placed on an agar pad (LB + 2% agar), and covered with a cover slip to prevent evaporation.

Cell preparation for growth in colonies was closely matched to the protocol for growing colonies to measure demographic noise in this work. Cultures were grown overnight with rotation at 37C. The next day, a $2\mu$L droplet of the saturated culture was placed on a plate with LB and 2% agar. The colony was grown at 37C for 1.5 days. Cells were picked from the edge of the colony and resuspended in LB, and vortexed to break up clumps. For imaging, a droplet of the resuspended culture was placed on an agar pad (LB + 2% agar), and covered with a cover slip to prevent evaporation.

Cells were imaged with phase contrast microscopy using a Nikon Eclipse Ti-E inverted microscope with a 40x, 0.65 NA phase contrast air objective (Nikon, Düsseldorf, Germany). Image segmentation was done with the software Morphometrics [16] and features that were incorrectly segmented were manually rejected. The circularity $C$ is defined as

$$C = \frac{l^2}{4\pi A} \tag{3}$$

where $l$ is the cell contour length, and $A$ is the cell area. The circularity of a circle is 1 and values larger than 1 correspond to more elongated cells.

#### 1.9 Single cell imaging with beads

In order to track the movement of the cells and beads at the single-cell resolution (Figure 1c and Figure S1c), we grew a colony following the procedure in the main experiments for demographic noise for 18-24 hours at 37C, then covered the colony with a BSA-coated coverslip. The sample was incubated at 32C with a H201-T Okolab incubator (Ottaviano, Italy) and a home-built incubation chamber for imaging. We imaged the cells and beads with an Olympus IX81 Inverted microscope with a 40x 1.3 NA oil objective (Olympus, Hamburg, Germany) in brightfield. Images were taken every 2-4 minutes for approximately 1 hour. To correct for focus drift due to evaporation, we used $\mu$Manager to move the z position of the stage at a constant rate and we also took a z stack of eleven $1\mu$m slices which were postprocessed to find the most in focus image. The beads could be identified from their different scattering properties compared to the cells.

#### 1.10 Gene enrichment analysis

GO and KEGG terms associated with at least 3 genes in the 191 randomly chosen strains were tested for the hypothesis that the weighted median MSD value of the knockouts associated with a given term was more extreme than that a randomly chosen subset of the same number of genes from all knockouts. For each term, random subsets of MSD values were drawn $10^5$ times. The fraction of times that the weighted median of the random subset was more extreme than that of the true values was recorded and the p-value was calculated as twice this fraction (due to only looking at the distribution on one side of the median). A Benjamini-Hochberg FDR correction [3] was applied to account for multiple testing, and terms that were significant at a 5% level were kept.

#### 1.11 Lasso regression

We used the Lasso regression implementation in the Python library scikit-learn [14] to determine the minimal set of traits that predict the MSD and the coefficients associated with those traits in a linear model. The data from each input phenotype is standardized to remove the mean and scale to unit variance. The cost for the regularization term is determined using 10-fold cross validation to be $\alpha = 0.015$.

#### 1.12 Partial correlation

The partial correlation was determined by first calculating the linear least squares regression coefficients between the trait of interest and all other traits, and between the bead MSD and all other traits except the trait of interest. Where errors for the trait of interest existed, they were used to weight the least squares fitting with a weight of $1/err^2$. The residuals were calculated for the trait of interest and the bead MSD. The partial correlation was calculated as the correlation between the residuals.

#### 1.13 Comparing establishment probability to bead MSD (Figure 4b)

For each genotype, at a particular chloramphenicol concentration, the replicate fitness (N=8) and establishment probability measurements (N=24) were randomly paired. A linear fit was performed using numpy.polyfit on the establishment probability as a function of fitness across all chloramphenicol concentrations and all initial mutant fractions for the fitness coefficient range from $s$ = -0.1 to $s$ = 0.5 since the colony collision assay is only valid for small fitness coefficients [10] (see Figure S13). The establishment probability was calculated using the fit parameters at the fitness coefficients $s$ = 0.05, 0.1, and 0.15. The error in the establishment probability was taken as half of the difference between the maximum and minimum values possible using combinations of fitted slope and intercept parameters that are one standard deviation away. We note that while the establishment probability and fitness are not expected to depend linearly on one another according to theory [12], the linear relationship is a convenient approximation for our data where we don't have sufficient signal to distinguish between more complex models.

#### 1.14 Statistical methods

For calculating the confidence interval on the percent difference in medians between the knockout and wild type demographic noise distribution, a modified block jackknife method was used. $10^5$ subsets of 32 KO MSD values (the same number as in the WT distribution) were drawn (without replacement in each subset) and their medians were calculated. The confidence interval was taken from the 2.5% to the 97.5% of the empirically measured subsampled median distribution.

### 2 Supplemental text

#### 2.1 Additional information on bead-based sparse lineage tracing method in colonies

Demographic noise in colonies has previously been quantified by measuring the number or shape of sectors in whole colonies [8] or tracking single cell lineage dynamics in a growing microcolony [7]. However, these methods require fluorescently labeled strains or time-intensive imaging and analysis which makes them impractical for screening large numbers of strains.

Our label-free method to measure demographic noise in microbial colonies allows us to screen a large number of strains and to minimize the probability of accumulating new mutations by avoiding genetic transformation. This is important as we aim to test the effect of single loss of function mutations on demographic noise, and additional mutations could potentially change the strength of demographic noise that is measured. The method also allows potential future study of microbes that are not genetically tractable where genetic transformations with fluorescent markers would not be possible.

As described in the main text and shown in Figure 1a, we use spherical fluorescent polystyrene tracer beads to sparsely track cell lineages in colonies and we use the wandering statistics of the bead trajectories to infer the strength of demographic noise. Because the beads are at a lower spatial density than cells, we are able to image with an air objective with a lower NA and a lower magnification than if we imaged single cells, and we can also capture a larger field of view. We can also track the beads with images that are taken less frequently because beads move fewer pixels at a lower magnification, and this allows us to image many colonies in parallel.

As a first approach to validate the method, we compared the measurement of MSD from sector boundaries, single cell trajectories, and bead trajectories for *E. coli* DH5$\alpha$ and *S. cerevisiae* W303 (Figure S1e), which are two species whose strengths of demographic have previously been compared [7, 8]. In all three methods, *E. coli* has a higher MSD than *S.*

*cerevisiae* as previously measured [7, 8]. The three methods span different length scales, with the single cell lineages spanning the shortest length scale, the bead tracks spanning an intermediate length scale, and the sector boundaries spanning the longest length scale. There is a continuous transition from the MSD of *E. coli* single cells to beads; however, there is a discontinuity between the transition from bead trajectories to sector boundaries. Additionally, there are discontinuities in the transition between the three methods for *S. cerevisiae*. The discontinuity between the bead trajectories and the sector boundaries may result from movement of the sector boundaries behind the colony front due to continued growth and realignment of cells. Another source of the discontinuities may come from the finite spatial resolution for the sector and bead images.

As a second approach to validate the method, we imaged the beads and single cells for the wild type strain BW25113 in the same field of view to determine if the bead trajectories were following the cell lineages. Visually, we see that the bead trajectories follow at least 1 cell lineage (traced backwards in time) at the colony front and behind the front (Figure 1c and Figure S1c). We unfortunately were not able to image for a longer than 1 hour because the images went out of focus. Additionally, we note that adding a coverslip on top of the colony, which was necessary for using a high NA oil immersion objective to get single cell resolution, may change the mechanical behavior of the cells.

As a third approach to validate that the method can measure demographic noise, we compared the bead trajectory MSD to the fraction of diversity preserved in a neutral fluorescent mixture after 1 day of growth (Figure 1d). As discussed in the main text, we find fitting an inverse square root relationship to the fraction of diversity preserved as a function of the MSD roughly matches the shape of the relationship. We emphasize that our goal is not get the most accurate measurement of demographic noise, but to screen a large number of strains to determine whether there is a significantly different distribution of demographic noise effects from single gene knockouts compared to the wild type. Thus, small deviations from the fit can still allow us to measure the distribution of demographic noise as long as the deviations are less than the width of the measured distribution.

As beads can fall behind the front, the bead trajectories combine the dynamics at the front and behind the front. Since prior work has shown that successful lineages only come from the first layer of cells at the colony front [7], we tested how closely the full bead trajectories matched the bead trajectory only while it was at the front. We separately measured the MSD of trajectories at the front (within 6$\mu$m of the front) and behind the front. Figure S2 shows for wild type BW25113 that trajectories at the front exhibit a similar MSD to the full trajectories, whereas trajectories behind the front exhibit a lower MSD. Thus, the MSD of the full trajectories captures the MSD of trajectories at the front.

#### 2.2 Determining the mean squared displacement window size

In order to be able to compare a single number for the MSD for different genotypes, we reported a summary statistic of the MSD at a window size of $L = 50$ $\mu$m which is interpolated or extrapolated from the power law fit to the MSD across all window sizes. This window size was chosen because it gave the best fit of fraction of diversity preserved (y) to MSD (x) to the relationship $y = a\sqrt{x}$ [9] amongst the window sizes 25 $\mu$m, 50 $\mu$m, 100 $\mu$m, and 1000 $\mu$m (see Figure S3). The fit at 25 $\mu$m, 50 $\mu$m, and 100 $\mu$m gave similar reduced chi-squared values, while the fit at 1000 $\mu$m had a much higher reduced chi-squared. The poor fit at high window sizes is likely due to the fact that few bead trajectories become that long, and the extrapolation becomes noisy. Note that while the reduced chi-squared values are generally high, we were not able to account for the errors in MSD in the reduced chi-squared calculation; taking those errors into account should reduce the discrepancies in the summary statistic describing the deviation between the fit and the data.

We also show the distribution demographic noise for the wild type and knockout strains using the MSD reported at different window sizes in Figure S4. At all window sizes except the largest window size of $L = 1000\mu$m (where the MSD also did not agree as well with the fraction of diversity preserved) we see that the wild type and knockout distributions are significantly different from one another to at least a level of 5% (Kolmogorov-Smirnov test).

#### 2.3 Sources of variation

We tested for variation across positions on a plate and across different plates. While all media for the experiments described in the main text (except single cell tracking experiments) were made on the same day, variation between plates can result from: the exact ratios of LB powder, agar, and water, the temperature of the liquid as it is poured into a plate, the tilt of the plate while drying, the age of the plate at the time that it is used, and other hidden experimental parameters. Figure S5a shows that the Pearson correlation coefficient for the same strain grown in the same position on different plates was 0.4-0.82 for randomly sampled knockout strains (DE1-4), 0.65-0.83 for non-randomly sampled knockout strains (DE5), and 0.61-0.9 for *mreB* and *mrdA* single point mutation strains. From these plots, we see unusually low correlation of plate DE3c with its replicates, and we decided to remove it from further analysis. Inspecting the colonies grown on plate DE3c showed that the colonies tended to be larger and less thick than those of the other replicate plates. Thus, we hypothesize that its discrepancy from its replicates may be due to higher plate moisture. Figure S5b shows that the Pearson correlation coefficient for strains grown in different positions on the same plate was 0.55-0.67. Thus, we observe comparable levels of variation due to differences in a strain's position on the plate and differences between plates.

We also tested for variation that may result due to the noisiness of averaging a finite number of bead trajectories for each colony. For the selected knockout strains, we randomly split the bead trajectories in each colony's field of view in half, and calculated the MSD separately for each set of trajectories. Figure S5c shows that the Pearson correlation coefficient between the two random sets of half of the trajectories is 0.78. Thus, we see that while finite track numbers can lead to variation between replicates, it does not play the largest role in determining variation between replicates, as differences between plates generally leads to more variation.

As described in the main text, in order to correct for variations across plates when measuring the distribution of demographic noise effects, we grew 8 wild type colonies on each plate with the randomly selected knockout strains, varying their positions on each plate. We normalized the measurements for each colony to the average wild type value on that plate for all measurements (bead MSD and phenotypic traits) to get a relative measurement. This effectively removes any systematic differences between plates but does not remove additional non-systematic variation from differences between plates, variation between positions, or variation from the number of tracks in each field of view.

#### 2.4 Beneficial sectors in monoculture colonies

We filtered out colonies with a noticeable beneficial sector in the field of view of the time-lapse from the analysis as these beneficial mutants may change the measured strength of demographic noise if grown to a large enough size. We identify beneficial sectors first by looking for bead trajectories that are strongly diverging in space, indicating faster growth than surrounding cells. Next, we verify this in the brightfield timelapse by looking for bulges at the colony front that expand over time. The majority of the 161 genotypes that are filtered out of the 352 randomly selected knockouts are due to two or more replicates having beneficial sectors.

To better understand if the number of colonies with beneficial sectors is surprising, we can estimate the expected number of de novo beneficial mutations from growth in the colony:

$$m_{b,est} = \mu_b p_{est} N \left( \frac{T}{\tau_{gen}} \right) \tag{4}$$

where $\mu_b$ is the beneficial mutation rate per genome per generation, $p_{est}$ is the establishment probability, $N$ is the number of cells (for the colony, it's the number of cells in the front layer that are actively dividing), $T$ is the total time, and $\tau_{gen}$ is the generation time. Using $\mu_b = 10^{-5}$, $p_{est} > 10^{-3}$ (Ref [7]), $N = 10^9$, $T/\tau_{gen} = 10$, then the expected number of established de novo beneficial mutations from growth in a single colony is at least $m_{b,est} > 100$, which means that we should not be surprised to see de novo beneficial mutations that have established in the colonies. We also note that the number of colonies identified with beneficial sectors from the imaging data is an underestimate of the true number of beneficial mutations in the entire colony because we are not able to identify weak beneficial mutations or mutations that arise later on and stay a small size and we also only image about 1/10 of the colony front.

An additional contribution to the number of beneficial sectors could be standing variation in the glycerol stock. This was tested by streaking single colonies from the glycerol stock, and picking colonies to inoculate the culture as a condition without standing variation. Figure S6 shows that cultures started from single colonies exhibited beneficial sectors as well as those started from glycerol stock. Thus the beneficial sectors that we see are likely a combination of both de novo mutations and standing variation from glycerol stock.

#### 2.5 Gene enrichment analysis

We asked whether there are gene categories that are associated with extreme changes to demographic noise. To do this, we compared the weighted median MSD of gene knockouts associated with a particular GO or KEGG term with randomly chosen MSDs from the knockout distribution.

From this analysis, we did not find any significant GO terms, and we only found one significant KEGG term that was enriched for higher MSD than the weighted median of the entire knockout distribution: ATP-binding cassette transporters. It is possible that expanding the dataset by measuring more genotypes or decreasing the amount of technical noise by performing more experimental replicates may result in more hits in the gene enrichment analysis. However, another possibility for the few gene enrichment hits is that the influence of phenotypic traits on demographic noise dominates over any differences between gene categories.

#### 2.6 Additional measurements of distribution of drift effects

In the main text we presented results for the distribution of demographic noise for a randomly selected set of 191 single gene knockout strains, in order to get a representative of the distribution for all single gene knockouts. To test for extreme differences in demographic noise, we also enriched for strains that we hypothesized to significantly change demographic noise. This included (1) selected single gene knockout strains that had altered biofilm forming ability in liquid culture [13] or different cell shapes [1], and different *E. coli* strain backgrounds, and (2) *mreB* and (3) *mrdA* single point mutant libraries that were enriched for cell shape differences [15].

Figure S7 shows the distributions of demographic noise of the selected single gene knockouts, the *mreB* point mutants, and the *mrdA* point mutants, compared with the wild type BW25113 and the randomly selected knockouts. Note that the *mreB* and *mrdA* strains are on the MG1655 strain background, but here we will compare them to the wild type BW25113 measurements because we only made 2 measurements of MG1655 which is not enough to compare distributions and we found that MG1655 has a similar strength of demographic noise to BW25113 (Figure 1e).

The tail at the lower end of the distributions for all three additionally tested sets of strains does not overlap with the gaussian fit to the wild type distribution nor do they completely overlap the grayed out region from the finite number of wild type measurements. The selected single gene knockout distribution and the *mreB* point mutants distribution are significantly different from that of the wild type using a two-sample Kolmogorov-Smirnov test ($p = 1.9 \times 10^{-10}$ for selected knockouts, $6 \times 10^{-5}$ for *mreB* point mutants), while the *mrdA* point mutant library is less significantly different from the wild type distribution ($p = 6.8 \times 10^{-2}$), most likely due to the fewer number of measurements.

The distributions of the strength of demographic noise for the selected single gene knockouts and the *mreB* point mutants are wider than that of the randomly selected knockouts. Some strains even have MSD close to zero. Thus, we see that the maximally allowed changes to demographic noise are much more extreme than we have sampled with our randomly selected strains. Sampling more strains will also lead to a more extreme knockout distribution. Similarly to what's observed for the randomly selected knockouts, many more mutants have a lower strength of demographic noise than a higher strength of demographic noise compared to the wild type.

#### 2.7 Comparison of cell shape between growth in liquid culture and as a colony

We used existing datasets for the cell shape of single gene knockout strains in the Keio collection [16, 6]. However, cells in these previous studies were grown in liquid culture, and it is unclear whether the cell shape could be different in colonies (the condition in this study). To determine whether these datasets would be applied to our experiments, we measured cell shape from growth in liquid culture and colonies for a subset of 12 strains (see Supplementary Methods).

Figure S8c shows a Pearson correlation coefficient between the mean circularity of single cells between the colony and liquid culture growth conditions to be 0.46. However, this is primarily driven by the large circularity of the genotype $\Delta gpmI$. Removing it from the measurement of the correlation coefficient gives a low correlation of -0.1 for the remaining genotypes. Thus, in interpreting the lack of correlation between cell shape and bead MSD in the single gene knockouts, we cannot rule out that it is because the cell shape is different in the colonies than the liquid culture condition of the previous datasets. We also note that the two previous datasets [16, 6] in liquid culture do not have any correlation with one another, further suggesting cell shape is sensitive to the precise growth conditions (Figure S8d). We note that the dataset from Ref. [16] did not reproduce the extreme cell shapes in the current images on the Keio collection database, which may be because the images have been changed online since their analysis.

#### 2.8 Non-neutral experiments

Figure S14a shows the fitness difference of the resistant and susceptible pairs in different strain backgrounds as measured by a colony collision assay (see Methods). Interestingly, different strain backgrounds exhibit different fitness coefficients despite having the same plasmids. This may be due to differences in plasmid copy number or epistatic effects between the plasmid and the rest of the genome. Studying the interaction between the strain background and fitness due to an antibiotic resistance gene on a plasmid would be an interesting avenue for future work.

We tested multiple different initial mutant fractions in order to be able to resolve individual sectors for different strain backgrounds and fitnesses. Figure S15b shows the establishment probabilities for three different initial fractions of the resistant type that are approximately $p_i = 0.002$, 0.005, and 0.02. Figure S14b shows the actually measured initial fractions for each of these genotypes using colony counting; the general trend of initial fractions matches the expectation, but there are offsets between strains. While the ordering of the establishment probabilities of the strains is similar across initial mutant fractions, the values themselves are slightly different, with slightly higher establishment probabilities for a lower initial mutant fractions. For better statistics, we used data from all initial mutant fractions to generate Figure 4b. The largest contribution to experimental error is in the measurement of the initial mutant fraction due to poisson sampling from counting a finite number of CFUs. Additional noise in the data may result from slightly different chloramphenicol concentrations between the agar plates used for the fitness measurements and the plates used for establishment probability measurements.

### 3 Supplementary figures

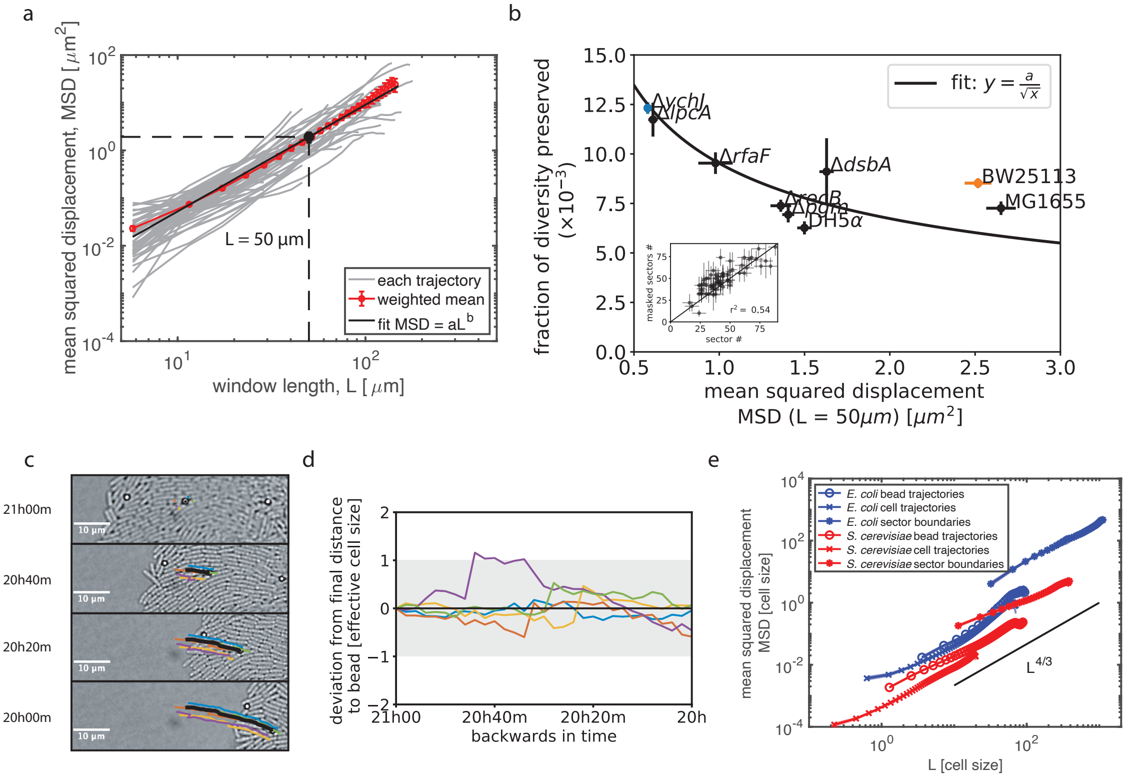


Figure S1: Bead-based sparse lineage tracing method. (a) Example of measurement of mean squared displacement (MSD) for a single colony (a single field of view). Gray lines show the MSD for single bead trajectories. The red line shows the average across all trajectories weighted by the inverse squared error of MSD for each trajectory at each window size. Error bars represent the standard error in the weighted mean. The black line shows the weighted least squares fit of the mean to a power law. The summary statistic MSD($L = 50$ $\mu$m) is calculated by interpolating the fit to $L = 50$ $\mu$m. (b) MSD(L $=$ 50 $\mu$m) compared to the fraction of diversity preserved when colonies are masked by the outline of the smallest colony to account for growth rate differences between colonies. Inset shows a comparison of the number of sectors counted with and without masking. (c) The trajectory of a single bead (black) behind the front and the lineages of cells neighboring it in the final timepoint (colors) traced backwards in time over 1 hour in wild type strain BW25113. (d) The deviation of the distance between the cell lineages and the bead from the final distance, backwards in time. Colors are the same as in (c). All cells neighboring the bead in the latest timepoint are neighboring the bead in the earliest timepoint, except for the yellow lineage (even though the yellow lineage does stay within a single cell width of its final distance to the bead). (e) *E. coli* has higher MSD than *S. cerevisiae* in all three methods for measuring demographic noise. Sector boundary mean squared displacement was measured according to [8]

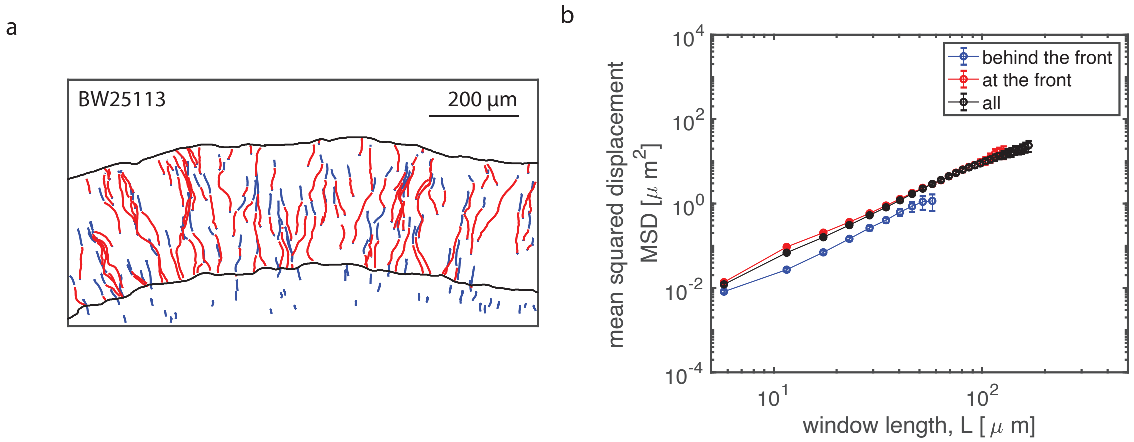


Figure S2: MSD of beads at and behind the front. (a) Trajectories of beads at the front (red, measured as within 6$\mu$m of the front) and behind the front (blue). (b) Beads at the front give similar MSD to that of overall bead trajectories.

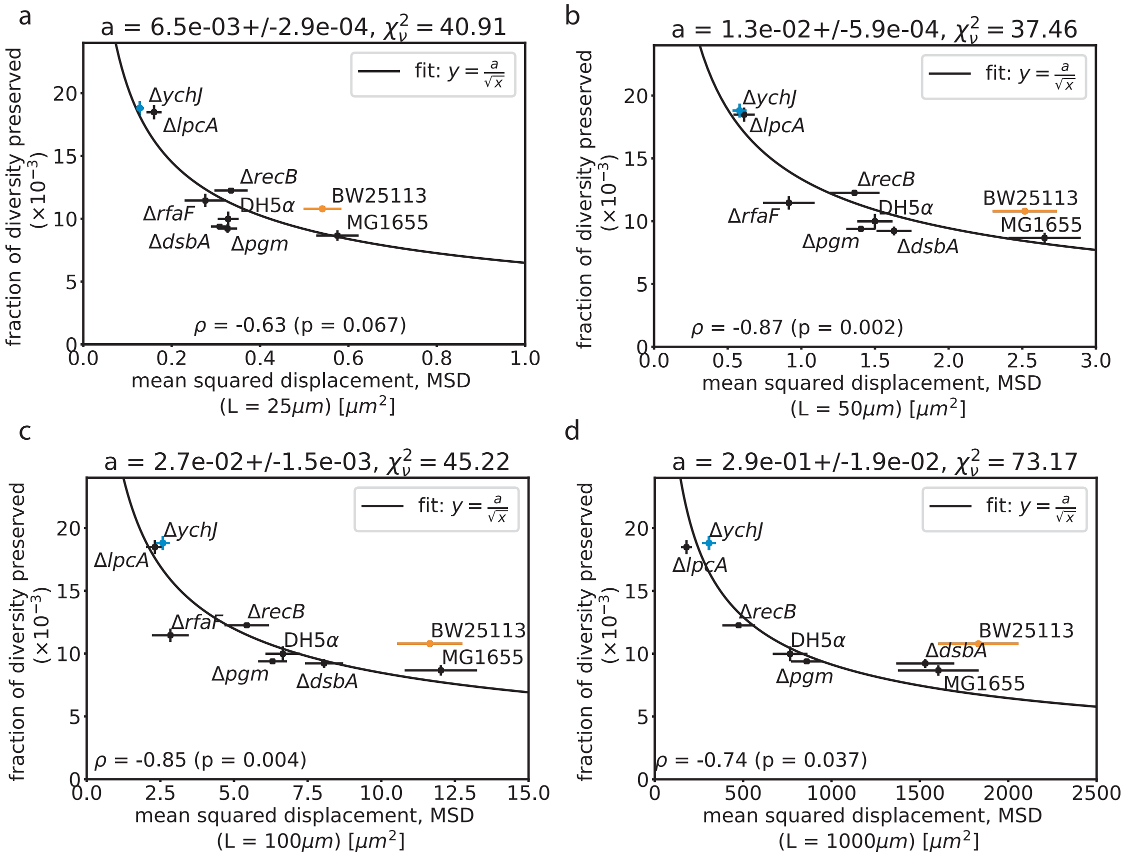


Figure S3: Selection of window size for the summary MSD statistic. Comparison of the fitted MSD value at different window sizes to the fraction of diversity preserved in a colony grown from a neutral mixture of two fluorescent strains for 24 hours. Error bars in MSD represent the standard error of the weighted mean (7-8 colonies) where weights come from uncertainties in the fit of MSD as a function of $L$ to a power law (see Methods) and error bars in the fraction of diversity preserved represent the standard error of the weighted mean (8 colonies) where weights come from uncertainties in counting the number of sectors . Fit to $y = ax^{-1/2}$, where $x$ is the MSD and $y$ is the fraction of diversity preserved, shows that the window size of $L = 50$ $\mu$m gives the lowest chi-squared value.

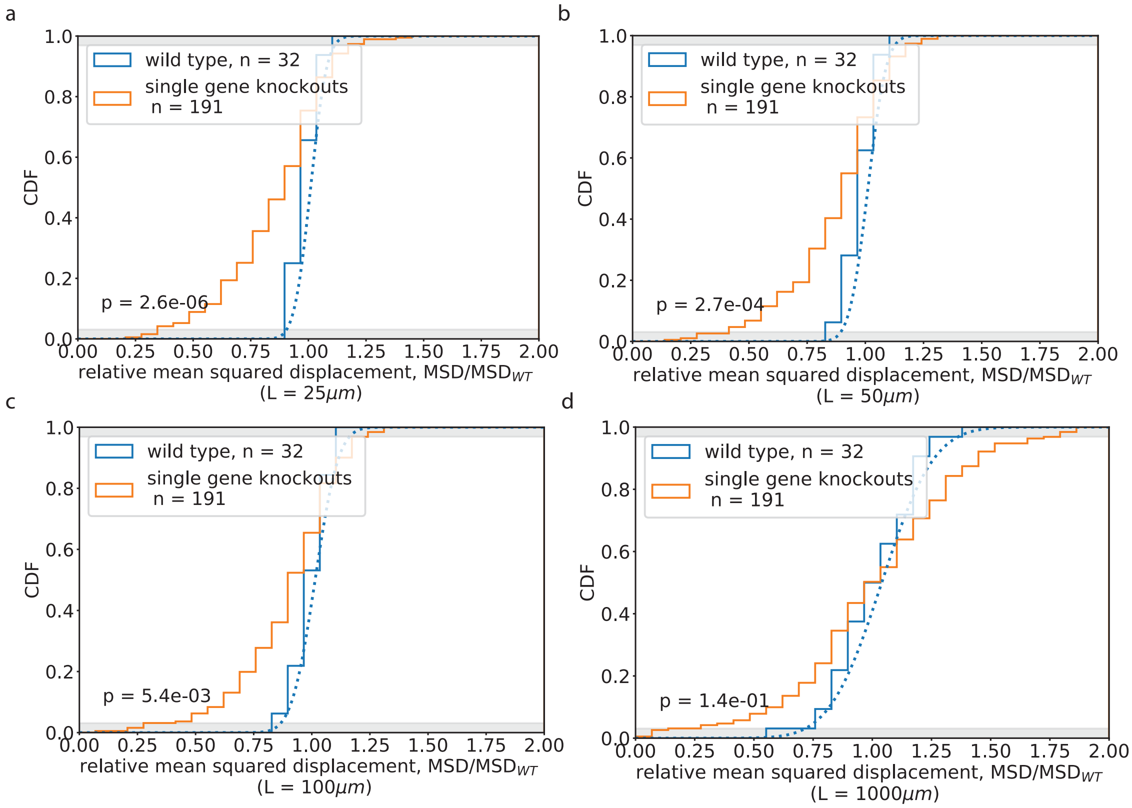


Figure S4: Distribution of demographic noise effects for the MSD reported at different window sizes. The blue dotted line shows a gaussian fit to the wild type distribution. The tails of the gaussian fit do not overlap with the tails of the knockout distribution. The gray shaded region shows [1/number of wild type measurements] and [1-(1/number of wild type measurements)], which is the limit of the resolution of the wild type distribution being compared to. p values show the probability that the wild type and knockout distributions are the same using a two sample Kolmogorov-Smirnov test. The knockout distribution is different from the wild type distribution to $p < 0.05$ for all window sizes except $L = 1000\mu$m.

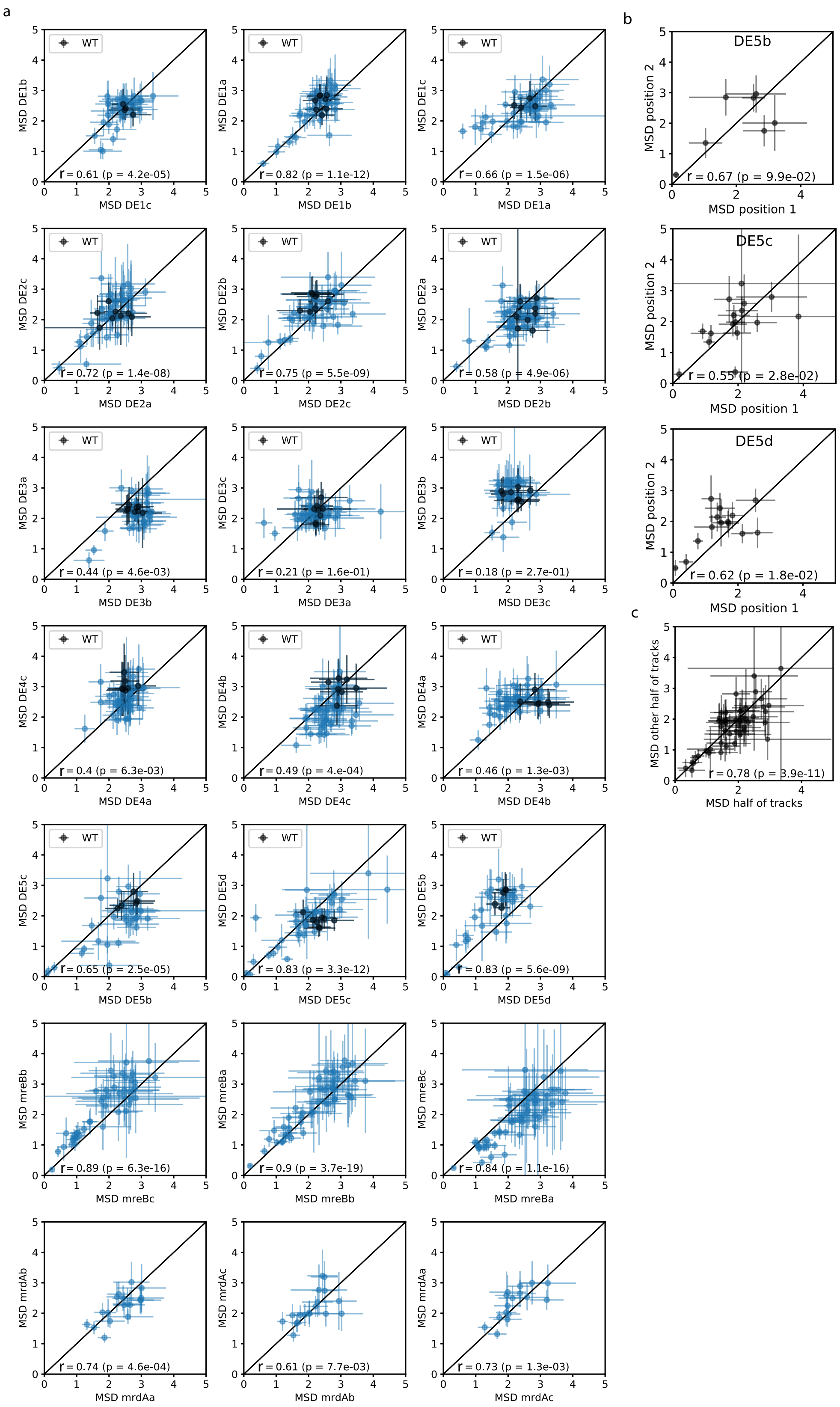
a
b
c
WT
MSD DE1b
MSD DE1c
r = 0.61 (p = 4.2e-05)
MSD DE1a
MSD DE1b
r = 0.82 (p = 1.1e-12)
MSD DE1c
MSD DE1a
r = 0.66 (p = 1.5e-06)
MSD DE2c
MSD DE2a
r = 0.72 (p = 1.4e-08)
MSD DE2b
MSD DE2c
r = 0.75 (p = 5.5e-09)
MSD DE2a
MSD DE2b
r = 0.58 (p = 4.9e-06)
MSD DE3a
MSD DE3b
r = 0.44 (p = 4.6e-03)
MSD DE3c
MSD DE3a
r = 0.21 (p = 1.6e-01)
MSD DE3b
MSD DE3c
r = 0.18 (p = 2.7e-01)
MSD DE4c
MSD DE4a
r = 0.4 (p = 6.3e-03)
MSD DE4b
MSD DE4c
r = 0.49 (p = 4.e-04)
MSD DE4a
MSD DE4b
r = 0.46 (p = 1.3e-03)
MSD DE5c
MSD DE5b
r = 0.65 (p = 2.5e-05)
MSD DE5d
MSD DE5c
r = 0.83 (p = 3.3e-12)
MSD DE5b
MSD DE5d
r = 0.83 (p = 5.6e-09)
MSD mreBb
MSD mreBc
r = 0.89 (p = 6.3e-16)
MSD mreBa
MSD mreBb
r = 0.9 (p = 3.7e-19)
MSD mreBc
MSD mreBa
r = 0.84 (p = 1.1e-16)
MSD mrdAb
MSD mrdAa
r = 0.74 (p = 4.6e-04)
MSD mrdAc
MSD mrdAb
r = 0.61 (p = 7.7e-03)
MSD mrdAa
MSD mrdAc
r = 0.73 (p = 1.3e-03)
DE5b
MSD position 2
MSD position 1
r = 0.67 (p = 9.9e-02)
DE5c
MSD position 2
MSD position 1
r = 0.55 (p = 2.8e-02)
DE5d
MSD position 2
MSD position 1
r = 0.62 (p = 1.8e-02)
MSD other half of tracks
MSD half of tracks
r = 0.78 (p = 3.9e-11)

Figure S5 *(preceding page)*: Sources of variation (a) Comparison of MSD for strains grown in the same position on different plates (r = 0.4-0.82 for randomly sampled knockout strains DE1-4, r = 0.65-0.83 for non-randomly sampled knockout strains DE5, and r = 0.61-0.9 for *mreB* and *mrdA* single point mutants). Wild type strains on each plate indicated in black. The plate DE3c exhibited low correlation with its replicates and was removed from further analyses. (b) Comparison of MSD for strains grown in different positions on the same plate (r = 0.55-0.67). (c) Comparison of MSD from randomly splitting the bead trajectories from a single colony in half and separately calculating MSD for each set of trajectories (r = 0.78).

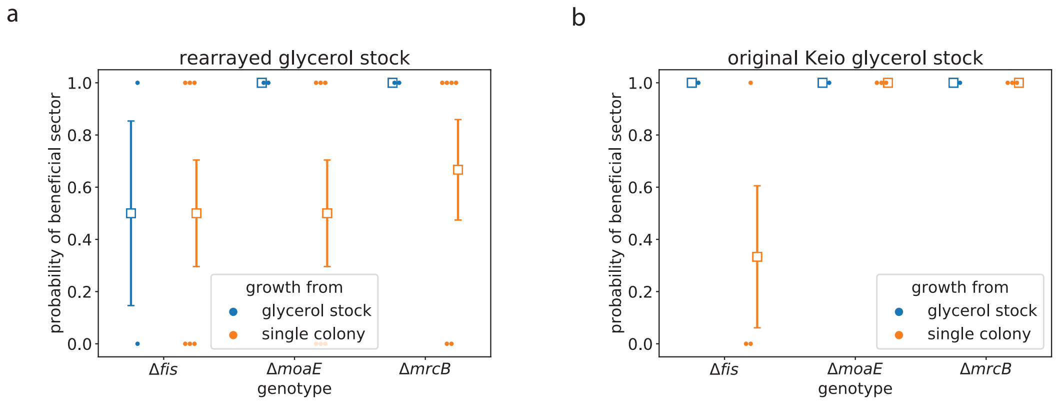


Figure S6: Testing for standing variation vs de novo mutations. Probability of seeing at least 1 beneficial sector in colonies grown from (a) the rearrayed glycerol stock or (b) the original Keio collection glycerol stock received from the National BioResource Project. Colonies were grown from liquid culture that was inoculated either directly by scraping off some glycerol stock (blue) or by first streaking the glycerol stock onto a plate, and picking from a single colony (orange). Otherwise, the experimental conditions were the same as that for measuring the distribution of demographic noise. Dots show individual colonies with none (0) or at least 1 beneficial sector (1) and squares show the average across colonies, which represents the probability of seeing a beneficial sector in a given condition. The error bars represent the standard deviation from binomial sampling. In all conditions, beneficial sectors were observed in at least one colony. This supports the hypothesis that beneficial mutations can arise de novo on the timescale of the experiment. Most likely, both beneficial mutations and standing variation from the glycerol stock contribute to the observation of beneficial sectors in the main experiment.

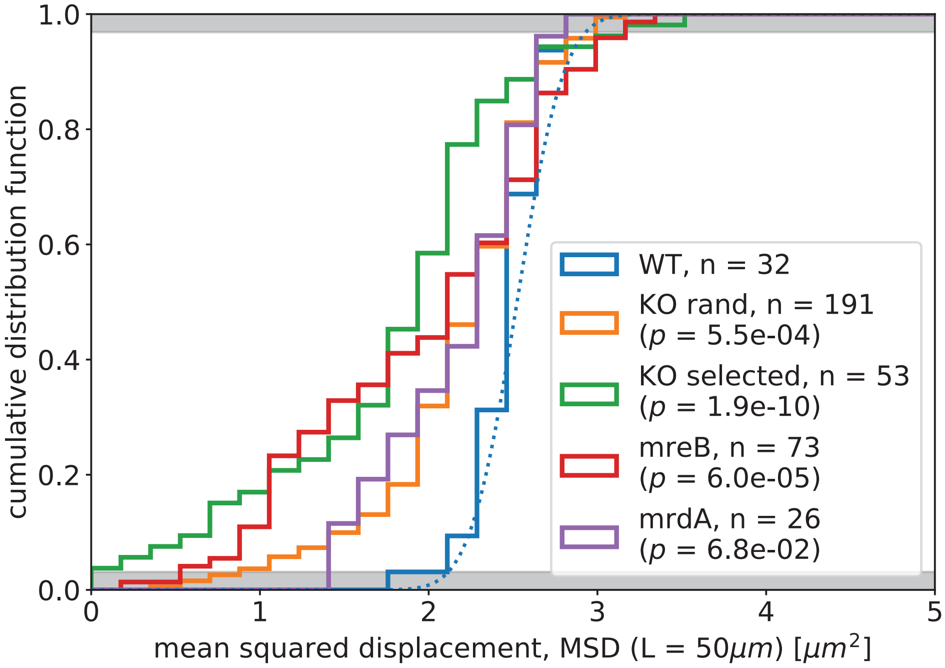


Figure S7: Additional measurements of demographic noise effects in non-randomly selected strains. (a) The distribution of strengths of demographic noise for the wild type BW25113 (WT), randomly selected single gene knockouts (KO rand), specifically selected single gene knockouts and strain backgrounds (KO selected), *mreB* single point mutants (*mreB*), and *mrdA* single point mutants (*mrdA*). All except *mrdA* single gene knockouts show a significantly different distribution of demographic noise effects compared to the WT to a significance level of $p <0.05$.

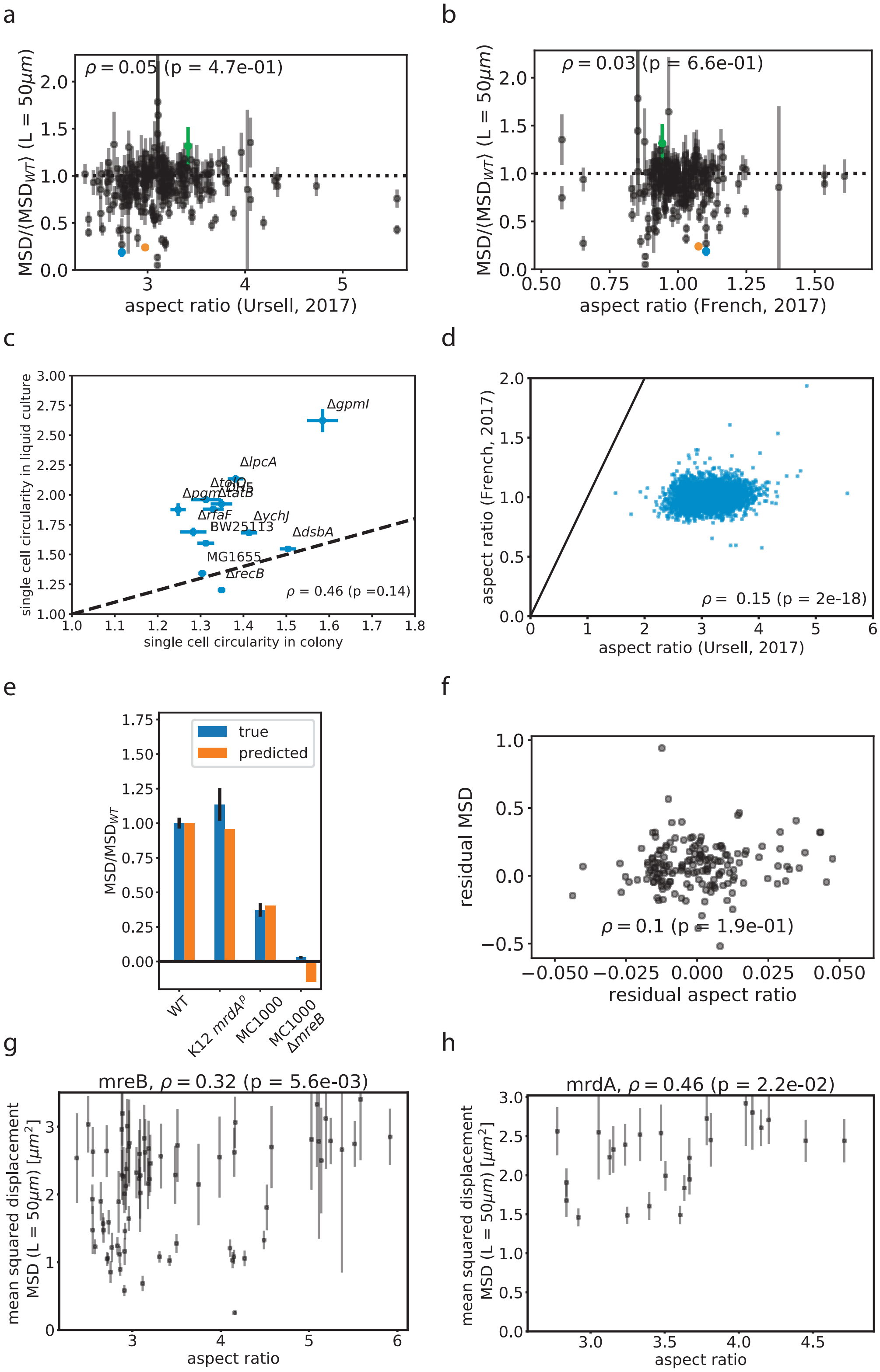
a
ρ = 0.05 (p = 4.7e-01)
MSD/⟨MSD_WT⟩ (L = 50μm)
aspect ratio (Ursell, 2017)
b
ρ = 0.03 (p = 6.6e-01)
MSD/⟨MSD_WT⟩ (L = 50μm)
aspect ratio (French, 2017)
c
ΔgpmI
ΔlpcA
ΔtolQ
Δpgm
ΔtatB
ΔrfaF
ΔychJ
BW25113
ΔdsbA
MG1655
ΔrecB
ρ = 0.46 (p =0.14)
single cell circularity in liquid culture
single cell circularity in colony
d
aspect ratio (French, 2017)
ρ = 0.15 (p = 2e-18)
aspect ratio (Ursell, 2017)
e
true
predicted
MSD/MSD_WT
WT
K12 mrdA^p
MC1000
MC1000 ΔmreB
f
residual MSD
ρ = 0.1 (p = 1.9e-01)
residual aspect ratio
g
mreB, ρ = 0.32 (p = 5.6e-03)
mean squared displacement MSD (L = 50μm) [μm²]
aspect ratio
h
mrdA, ρ = 0.46 (p = 2.2e-02)
mean squared displacement MSD (L = 50μm) [μm²]
aspect ratio

Figure S8 *(preceding page)*: Tests of correlation between cell shape and demographic noise. The strength of demographic noise is uncorrelated with cell aspect ratio in the 191 randomly sampled strains from the Keio collection and 34 selected strains from the Keio collection with cell shape data taken from (a) Ursell et al [16] and (b) French et al [6]. (c) The circularity (see Supplementary methods) of cells grown in liquid culture compared to those grown in a colony. Dashed black line depicts 1:1 relationship. The Pearson correlation coefficient is 0.46 with all genotypes, but only -0.1 when excluding $\Delta gpmI$. The error bars represent the standard error of the mean across the 50-100 cells measured per genotype. (d) Comparing cell shape data from Ursell et al and French et al across all single gene knockouts in the Keio collection. (e) Relative bead trajectory MSD in cell shape mutants and those predicted by the best fit Lasso regression model from the main text which primarily includes colony-level traits. (f) Partial correlation of cell aspect ratio when controlling for all other phenotypes for the random set of single gene knockouts from the Keio collection. Correlation of demographic noise to single cell aspect ratio in *mreB* (g) and *mrdA* (h) single point mutants is higher than that seen for the single gene knockouts, possibly because these strains were enriched for cell shape differences. Strains and data from Ref [15].

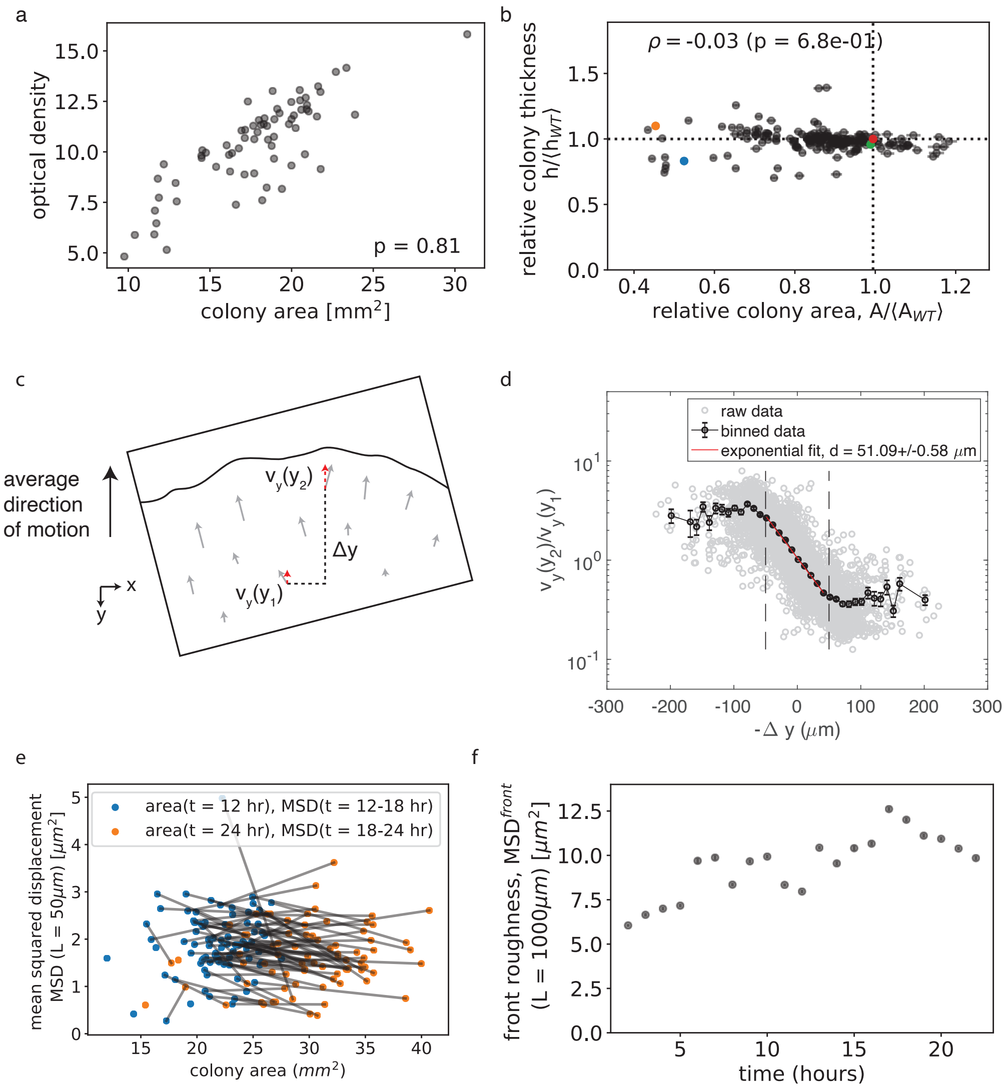

a
optical density
colony area [mm$^2$]
p = 0.81
b
$\rho$ = -0.03 (p = 6.8e-01)
relative colony thickness h/⟨h$_{WT}$⟩
relative colony area, A/⟨A$_{WT}$⟩
c
average direction of motion
$v_y(y_2)$
$v_y(y_1)$
$\Delta$y
x
y
d
raw data
binned data
exponential fit, d = 51.09+/-0.58 $\mu$m
$v_y(y_2)/v_y(y_1)$
-$\Delta$ y ($\mu$m)
e
area(t = 12 hr), MSD(t = 12-18 hr)
area(t = 24 hr), MSD(t = 18-24 hr)
mean squared displacement MSD (L = 50$\mu m$) [$\mu m^2$]
colony area ($mm^2$)
f
front roughness, MSD$^{front}$ (L = 1000$\mu m$) [$\mu m^2$]
time (hours)

Figure S9 *(preceding page)*: Additional tests of phenotypic traits (a) Colony area is a highly correlated measure of the optical density of the colony resuspended in liquid, which gives the total (alive and dead) biomass. (b) Colony thickness (see Methods) is not correlated with colony area, showing that area is a good measure of growth, rather than just spreading on the surface of the plate. (c) Schematic of measurement growth layer depth using bead displacements from a single colony's field of view. For a consecutive pair of timepoints, $\Delta y$ gives the distance between a pair of beads along the average direction of motion, and and $v_y(y_1)$ and $v_y(y_2)$ give the bead displacements projected onto the average direction of motion. (d) Gray points show measurements for all pairs of beads across all pairs of consecutive time points. Black points give binned measurements. Error bars represent standard error of the mean. The binned points roughly followed an exponential decay function between $\Delta y =$ -50$\mu$m and $\Delta y = 50\mu$m, and we fit to an exponential decay function within this range, as measurements of bead pairs that are farther away may be dominated by noise. The decay length, $d$, is extracted from the fit and taken as the growth layer depth. (e) The MSD for bead trajectories from 12-18 hours as a function of the area of the colony at 12 hours (blue points) and the MSD for bead trajectories from 18-24 hours as a function of the area of the colony at 24 hours (orange points). The same colony over time is connected by a gray line. Bead trajectory MSD mostly decreases with increasing colony area over time. Thus, the positive correlation between colony area and MSD seen across genotypes cannot be explained by changes in the bead trajectories over time. (f) Colony front roughness (see Methods) for *E. coli* DH5$\alpha$ over time. Colony front roughness increases initially but levels off around 19 hours. Since we measure the colony front roughness at 24 hours, we expect to have passed the time when the behavior of the front roughness is transient. Note that this measurement was done on a colony grown in a 10cm diameter petri dish rather than an omniplate, and the colony may have access to a different total amount of nutrients. However, while the growth condition does not exactly match that in the experiment, we used this data to better understand the approximate timescales of when front roughness saturates in time.

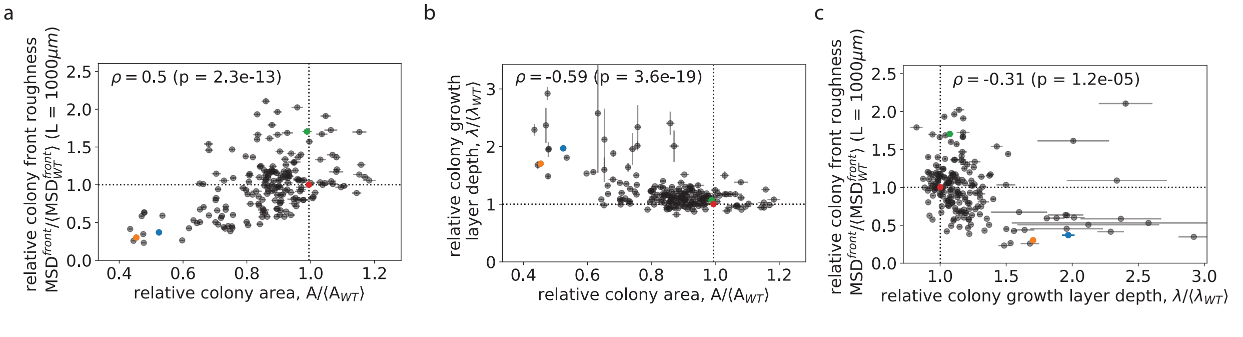


Figure S10: Correlation of colony traits with one another.

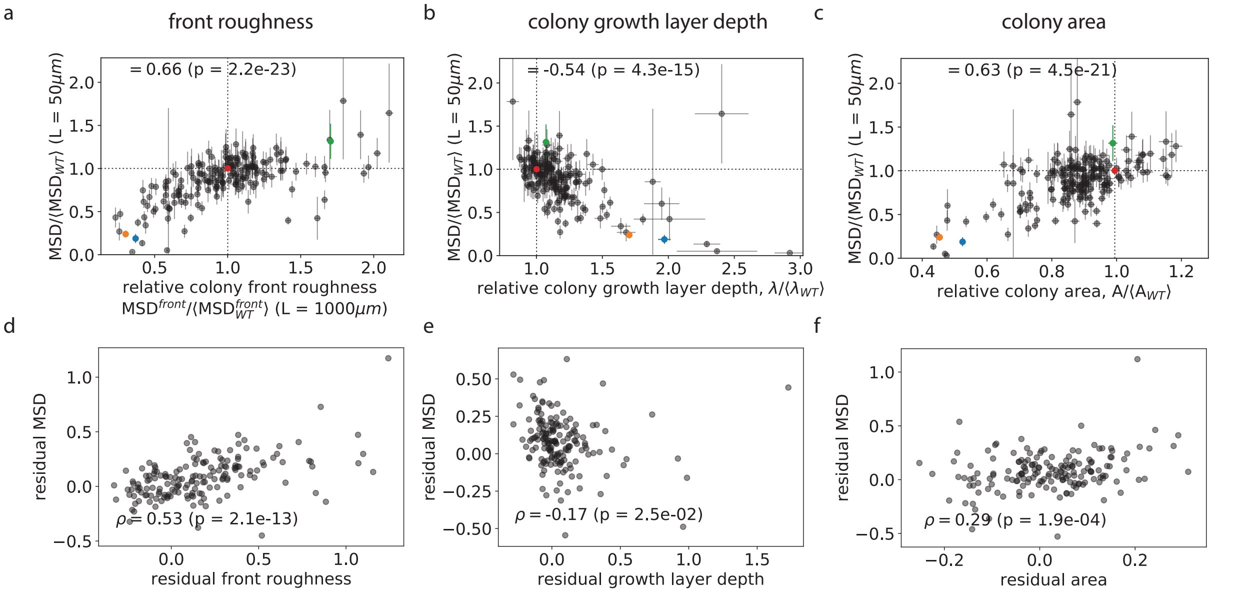


Figure S11: Correlation between the bead MSD and (a) front roughness, (b) growth layer depth, or (c) colony area (same as in Figure 3). Below each plot is the partial correlation between the bead MSD and each phenotypic trait (after controlling for correlations with all other traits).

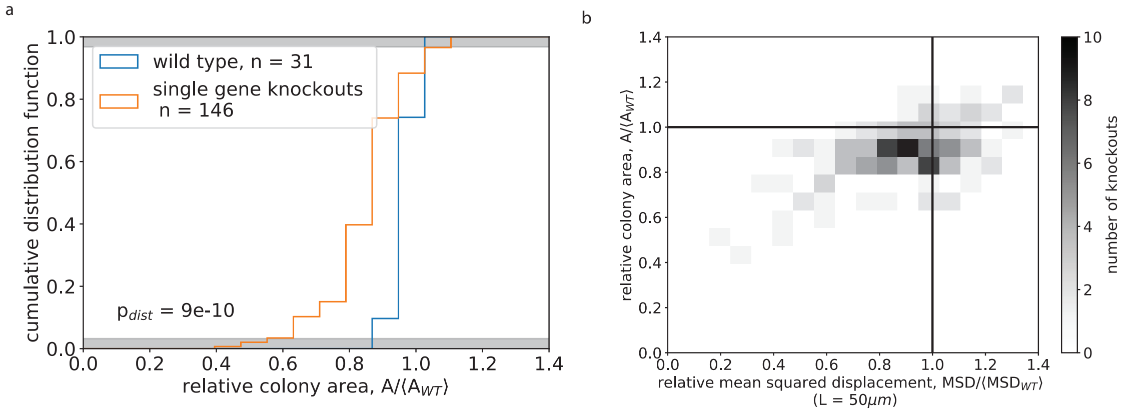


Figure S12: Joint distribution of demographic noise and colony fitness as measured by colony area. (a) The distributions of colony areas for the wild type BW25113 strain and the randomly selected single gene knockout strains. The distributions are significantly different to $p = 9 \times 10^{-10}$ as measured by a two-sample Kolmogorov-Smirnov test. (b) The joint distribution of demographic noise (as measured by bead MSD) and fitness (as measured by colony area) effects. The strength of demographic noise and fitness are correlated suggesting a tradeoff between fitness and demographic noise, with the median of both slightly below that of the wild type (black lines).

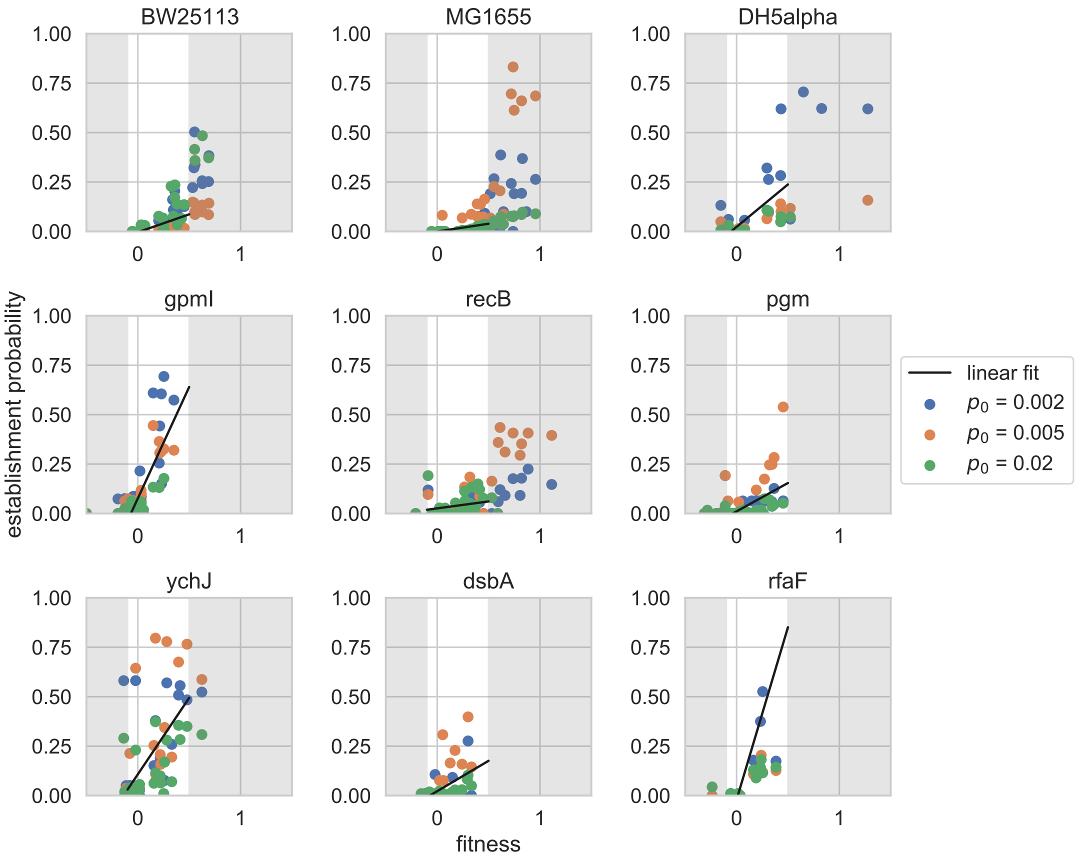


Figure S13: The establishment probability as a function of the fitness coefficient of a chloramphenicol resistant mutant for 9 selected strain backgrounds. Points that fall within fitness coefficients $-0.1 < s < 0.5$, where the colony collision assay is valid, are fit linearly (black line).

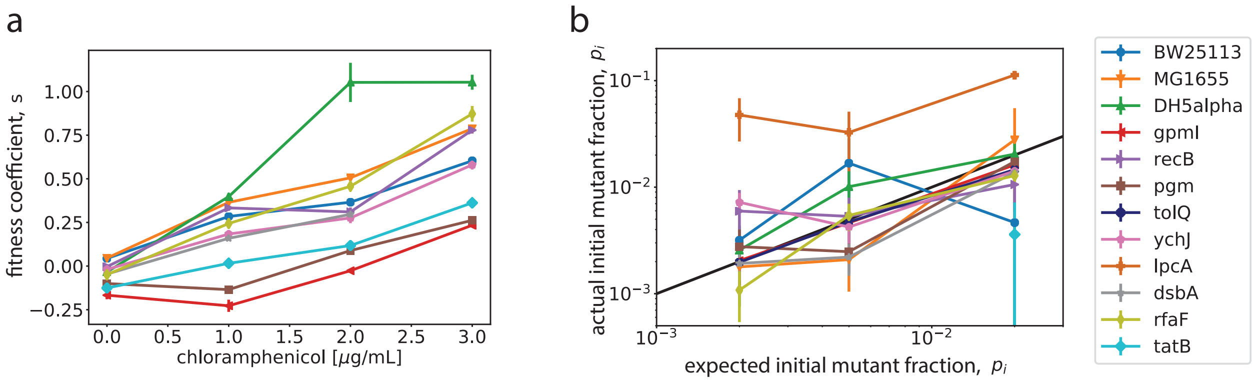


Figure S14: (a) The fitness coefficient ($s$) of a chloramphenicol resistant strain compared to chloramphenicol sensitive strain across strain backgrounds as measured on plates using a colony collision assay (Methods). (b) Expected and actual initial fraction of the resistant strain. Black line shows 1:1 relationship. $\Delta tatB$ had a large error in the measurement of the initial fraction of the mutant type from counting CFUs and was removed from Figure 4 in the main text and Figure S15.

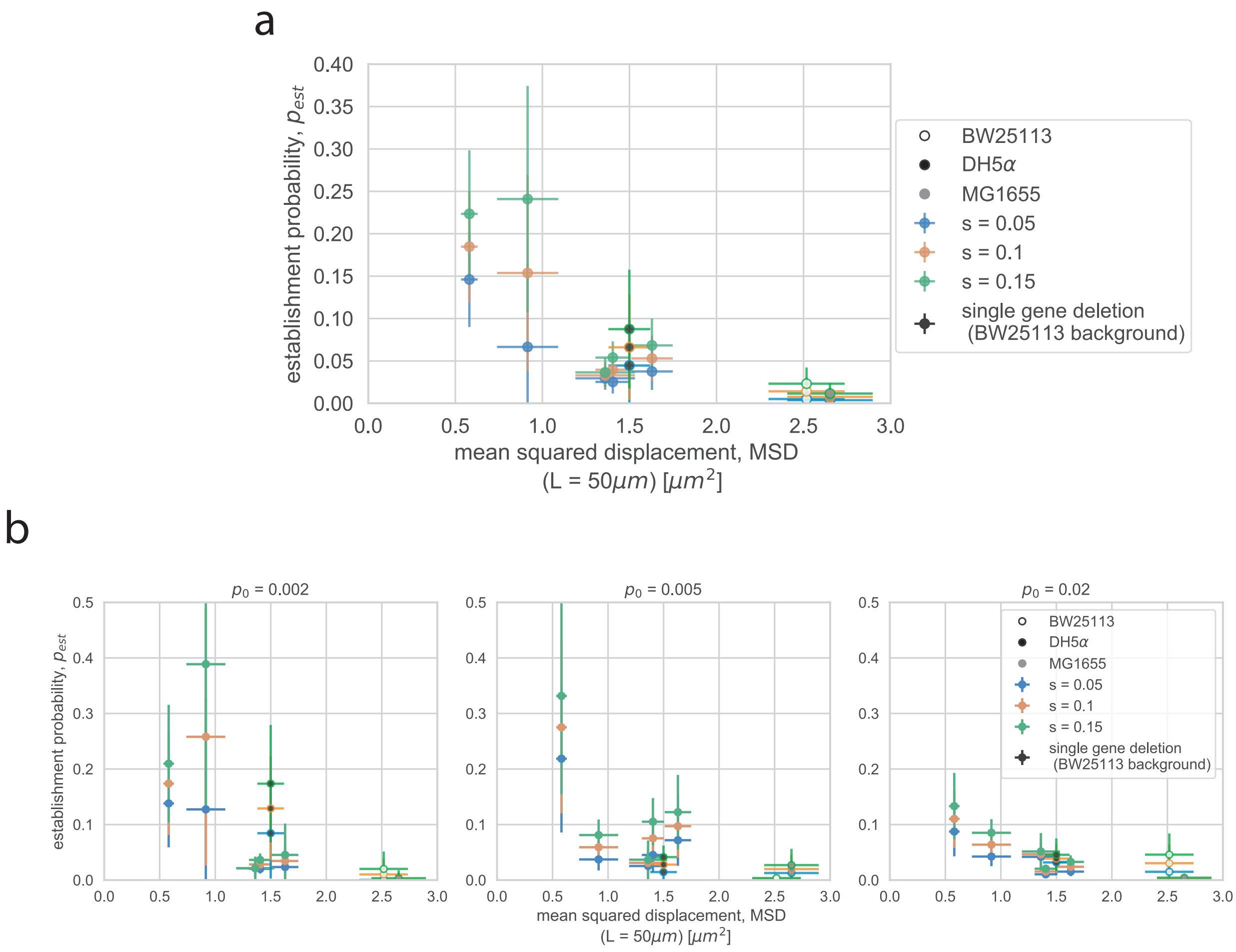


Figure S15: Fitted establishment probability at three different fitness coefficients from fitting (a) all initial resistant fractions together and (b) each initial resistant fraction separately as a function of bead trajectory MSD for 5 selected single gene deletion strains and 3 wild type strains. Error bars in the establishment probability represent linear fitting error (see Supplementary methods) and and error bars in MSD represent the standard error of the weighted mean (N = 7-8, see Methods).

### 4 Additional supplementary files

Supplementary movie 1: Movie of fluorescent beads being pushed by growing colony of *E. coli* DH5$\alpha$.

Type of file: movie

Supplementary table 1: List of *E. coli* single gene deletion strains used from Keio collection and plate positions of all strains tested

Type of file: table

Supplementary table 2: List of *mreB* and *mrdA* single point mutant strains used

Type of file: table

### References

[1] National bioresource project keio collection. `https://shigen.nig.ac.jp/ecoli/strain/resource/keioCollection/list`.

[2] Tomoya Baba, Takeshi Ara, Miki Hasegawa, Yuki Takai, Yoshiko Okumura, Miki Baba, Kirill A Datsenko, Masaru Tomita, Barry L Wanner, and Hirotada Mori. Construction of Escherichia coli K-12 in-frame, single-gene knockout mutants: the Keio collection. *Molecular Systems Biology*, 2(1):2006.0008, January 2006.

[3] Yoav Benjamini and Yosef Hochberg. Controlling the False Discovery Rate: A Practical and Powerful Approach to Multiple Testing. *Journal of the Royal Statistical Society: Series B (Methodological)*, 57(1):289–300, 1995. _eprint: https://rss.onlinelibrary.wiley.com/doi/pdf/10.1111/j.2517-6161.1995.tb02031.x.

[4] Marie-Cécilia Duvernoy, Thierry Mora, Maxime Ardré, Vincent Croquette, David Bensimon, Catherine Quilliet, Jean-Marc Ghigo, Martial Balland, Christophe Beloin, Sigolène

Lecuyer, et al. Asymmetric adhesion of rod-shaped bacteria controls microcolony morphogenesis. *Nature communications*, 9(1):1–10, 2018.
